# A clinal polymorphism in the insulin signaling transcription factor *foxo* contributes to life-history adaptation in *Drosophila*

**DOI:** 10.1101/473231

**Authors:** Esra Durmaz, Subhash Rajpurohit, Nicolas Betancourt, Daniel K. Fabian, Martin Kapun, Paul Schmidt, Thomas Flatt

## Abstract

A fundamental aim of adaptation genomics is to identify polymorphisms that underpin variation in fitness traits. In *D. melanogaster* latitudinal life-history clines exist on multiple continents and make an excellent system for dissecting the genetics of adaptation. We have previously identified numerous clinal SNPs in insulin/insulin-like growth factor signaling (IIS), a pathway known from mutant studies to affect life history. However, the effects of natural variants in this pathway remain poorly understood. Here we investigate how two clinal alternative alleles at *foxo*, a transcriptional effector of IIS, affect fitness components (viability, size, starvation resistance, fat content). We assessed this polymorphism from the North American cline by reconstituting outbred populations, fixed for either the low- or high-latitude allele, from inbred DGRP lines. Since diet and temperature modulate IIS, we phenotyped alleles across two temperatures (18°C, 25°C) and two diets differing in sugar source and content. Consistent with clinal expectations, the high-latitude allele conferred larger body size and reduced wing loading. Alleles also differed in starvation resistance and expression of *InR*, a transcriptional target of FOXO. Allelic reaction norms were mostly parallel, with few GxE interactions. Together, our results suggest that variation in IIS makes a major contribution to clinal life-history adaptation.

## Introduction

Much has been learned about the genetics of fitness traits (e.g., size, lifespan), mainly from studies of large-effect mutants and transgenes in yeast, *C. elegans*, *Drosophila* and the mouse (Finch and Rose 1995; Oldham and Hafen 2003; Tatar et al. 2003; Fielenbach and Antebi 2008; Kenyon 2010; Flatt and Partridge 2018), but loci identified in such laboratory analyses do not necessarily harbor segregating alleles that would contribute to genetic variance for traits in natural populations (Flatt 2004; Flatt and Schmidt 2009; Vonesch et al. 2016; Birney 2016; Fabian et al. 2018). In particular, the identity and presumably subtle effects of naturally occurring life-history polymorphisms are poorly known (Flatt and Schmidt 2009; Paaby and Schmidt 2009; Flatt and Heyland 2011). While adaptation genomics can in principle quite readily identify such candidate polymorphisms, a major – but rarely accomplished – objective is to experimentally validate these candidates as genic targets of selection (Barrett and Hoekstra 2011; Turner 2014; Flatt 2016; Siddiq et al. 2017). Thus, with a few exceptions, examples of causative life-history variants remain rare (Schmidt et al. 2008; McKechnie et al. 2010; Paaby et al. 2010; Jones et al. 2012; Johnston et al. 2013; Méndez-Vigo et al. 2013; Paaby et al. 2014; Barson et al. 2015; Catalán et al. 2016; reviewed in Mackay et al. 2009; Barrett and Hoekstra 2011).

Despite conceptual and methodological limitations of the so-called quantitative trait nucleotide (QTN) program (Rockman 2012), the identification of life-history polymorphisms allows addressing fundamental questions about the genetic basis of adaptation, including: (1) Which pathways and molecular functions underpin variation in fitness-related traits? (2) Are these mechanisms evolutionarily conserved? (3) What are the phenotypic effects of naturally segregating life-history variants? (4) What is the molecular nature of life-history epistasis, pleiotropy and trade-offs? (5) Do life-history polymorphisms mediate plasticity and how? (6) Is the genetic basis of evolutionary changes in life history ‘predictable’, i.e. relying on variation in the same pathways or genes? Or do life-history traits evolve unpredictably, i.e. via different pathways or loci, in different contexts?

A powerful model for dissecting the genetics of life-history adaptation is the vinegar fly *Drosophila melanogaster*, a species of sub-Saharan African origin, which has migrated out of Africa ~15,000 years ago and subsequently colonized the rest of the world (David and Bocquet 1975; David and Capy 1988; de Jong and Bochdanovits 2003; Hoffmann and Weeks 2007; Adrion et al. 2015). During the colonization of new climate zones, this ancestrally tropical insect has undergone a series of life-history adaptations to temperate, seasonal habitats (David and Capy 1988; de Jong and Bochdanovits 2003; Paaby and Schmidt 2009). This is particularly evident in the case of clines, i.e. directional patterns of phenotypic or genetic change across environmental gradients. Many studies have documented patterns of latitudinal differentiation among *D. melanogaster* populations that are presumably driven by spatially varying selection, for example along the North American and Australian east coasts, with the corresponding clines spanning subtropical/tropical and temperate habitats (de Jong and Bochdanovits 2003; Schmidt et al. 2005a, 2005b; Hoffmann and Weeks 2007; Schmidt and Paaby 2008; Kolaczkowski et al. 2011; Fabian et al. 2012; Adrion et al. 2015; Cogni et al. 2017). Clinal trait differentiation has been found, for instance, for body size, fecundity, reproductive dormancy, stress resistance and lifespan, typically in a parallel fashion on multiple continents, suggesting that these patterns are likely adaptive (Coyne and Beecham 1987; Schmidt et al. 2000; Weeks et al. 2002; de Jong and Bochdanovits 2003; Schmidt et al. 2005a, 2005b; Hoffmann and Weeks 2007; Schmidt and Paaby 2008; Adrion et al. 2015; Fabian et al. 2015; Kapun et al. 2016a).

To begin to identify the genetic basis of life-history clines in *D. melanogaster*, we have previously performed genome-wide analyses of latitudinal differentiation along the North American cline (Fabian et al. 2012; Kapun et al. 2016b) (also see Turner et al. 2008; Bergland et al. 2014; Reinhardt et al. 2014; Machado et al. 2018). Our analysis based on SNP *F*_ST_ outliers uncovered pervasive genome-wide patterns of clinality, with hundreds of clinal SNPs mapping to loci involved in the insulin/insulin-like growth factor signaling (IIS)/target of rapamycin (TOR), ecdysone, torso, EGFR, TGFβ/BMP, JAK/STAT, lipid metabolism, immunity and circadian rhythm pathways (Fabian et al. 2012). Many of the identified variants also exhibit parallel differentiation in Australia (Fabian et al. 2012; Kapun et al. 2016b; also cf. Kolaczkowski et al. 2011; Reinhardt et al. 2014; Machado et al. 2016), thereby strengthening the case for clinal adaptation. However, while many clinal variants might be shaped by selection, some of the observed differentiation might be due to non-adaptive factors, including population structure, demography, admixture or hitchhiking with causative sites (Endler 1977; Duchen et al. 2013; Kao et al. 2015; Bergland et al. 2016). Identifying adaptive clinal variants as targets of selection thus requires comparing clinal patterns against neutral expectations and – optimally – functional genetic testing (Barrett and Hoekstra 2011; Kapun et al. 2016a, 2016b; Flatt 2016). To date, however, functional analyses of clinal polymorphisms that are potentially subject to spatially varying selection remain scarce (for some exceptions see e.g. Schmidt et al. 2008; McKechnie et al. 2010; Paaby et al. 2010; Lee et al. 2013; Paaby et al. 2014; Kapun et al. 2016a; Durmaz et al. 2018; Svetec et al. 2018).

Interestingly, many of the pathways that harbor clinal loci are known from molecular studies to be implicated in the physiological regulation of life history in organisms such as *C. elegans*, *Drosophila* and the mouse (see Tatar et al. 2003; Fielenbach and Antebi 2008; Flatt and Heyland 2011; Flatt et al. 2013; and references therein). In particular, we found strongly clinal SNPs in multiple components of the IIS/TOR pathway, including SNPs in *Drosophila insulin-like peptide* genes *dilp3* and *dilp5*, *insulin-like receptor* (*InR*), *phosphatidyl-inositol-4,5-bis-phosphate 3-kinase* (*Pi3K*), forkhead box-O transcription factor *foxo*, the *foxo* regulator *14-3-3ε*, *target of brain insulin* (*tobi*)*, tuberous sclerosis complex 1* (*Tsc1*), and *target of rapamycin* (*Tor*) (Fig. 1; Fabian et al. 2012; Kapun et al. 2016b). This pattern is compelling since loss-of-function mutations in the IIS/TOR pathway have major, evolutionarily conserved effects on growth, size, reproduction, lifespan and stress resistance in *Drosophila*, *C. elegans*, and the mouse (Kenyon et al. 1993; Gems et al. 1998; Böhni et al. 1999; Brogiolo et al. 2001; Tatar and Yin 2001; Clancy et al. 2001; Kenyon 2001; Oldham et al. 2002; Oldham and Hafen 2003; Holzenberger et al. 2003; Tatar et al. 2001; Partridge et al. 2005).

**Figure 1.**
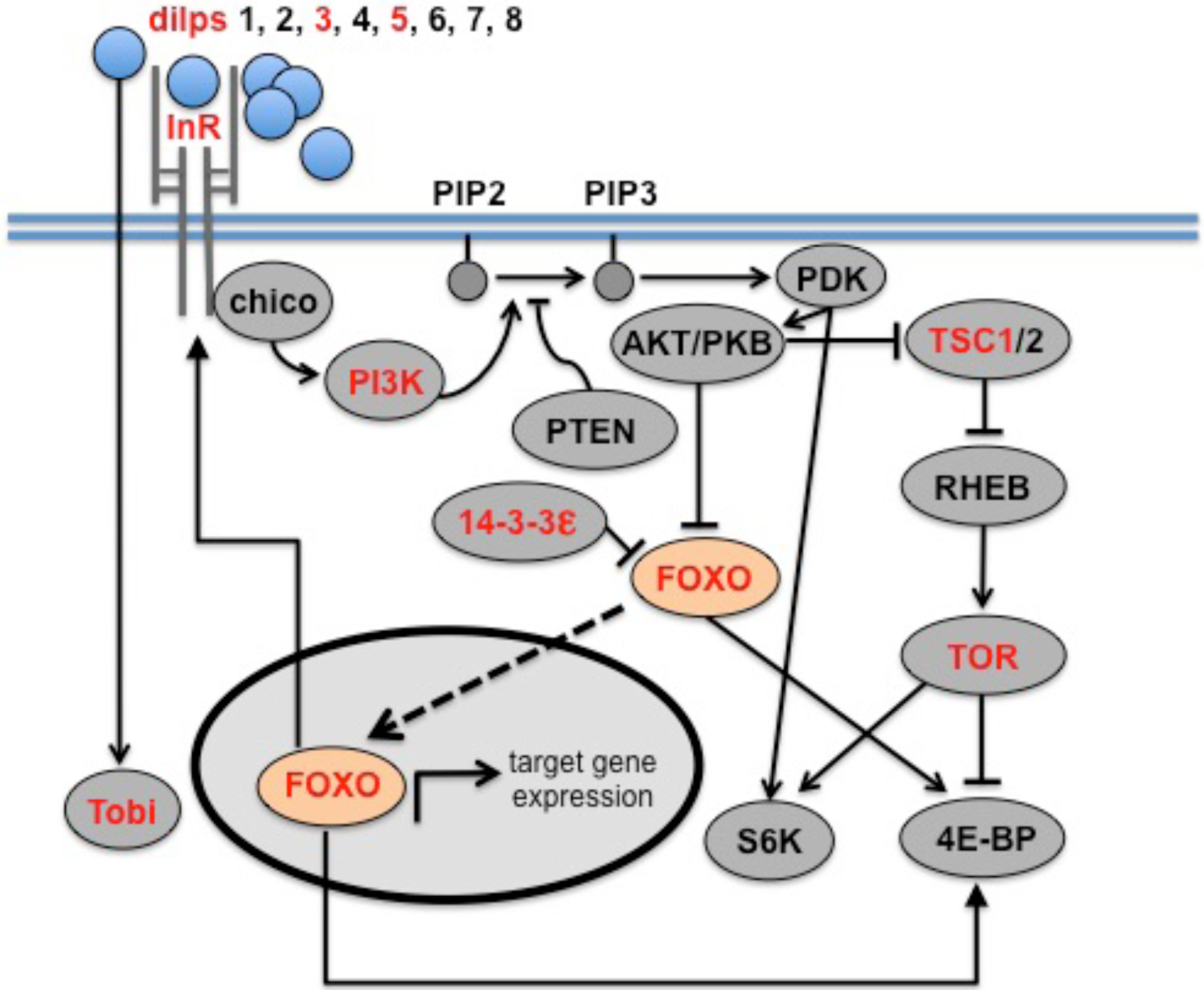
Clinal candidates in the insulin/TOR signaling pathway. Overview of the insulin/insulin-like growth factor signaling (IIS)/target of rapamycin (TOR) pathway in *Drosophila melanogaster* (Oldham and Hafen 2003; Giannakou and Partridge 2007; Teleman 2010). Genes that harbor strongly clinally varying SNPs across latitude, identified by Fabian et al. (2012), are highlighted in red; arrows indicate activation and bar-ended lines represent inhibitory effects. In response to nutrients, IIS is activated by binding of ligands, called *Drosophila* insulin-like peptides (dilps 1-8), to the insulin-like receptor (InR) at the cell membrane. Inside the cell, signaling is transduced by an insulin receptor substrate (IRS) protein called chico. This activates phosphoinositide-3-kinase (PI3K) which converts phosphatidylinositol (3,4)-bisphosphate (PIP2) into phosphatidylinositol (3,4,5)-trisphosphate (PIP3). In turn, PIP3 stimulates pyruvate dehydrogenase kinase (PDK) and activates protein kinase B (AKT/PKB). The action of PI3K is antagonized by phosphatase and tensin homologue (PTEN) which converts PIP3 back to PIP2. AKT/PKB suppresses the forkhead (FKH) box O transcription factor FOXO by phosphorylating it; upon reduced IIS, FOXO becomes dephosphorylated and moves into the nucleus where it regulates the expression of hundreds of target genes. Target genes of FOXO include *InR*, controlled via a transcriptional feedback loop, and *initiation factor 4E-binding protein* (*4E-BP*); another target gene of IIS is *target of brain insulin* (*Tobi*), which encodes a glucosidase, but the details of its regulation remain poorly understood. FOXO is antagonized by 14-3-3ε. AKT/PKB antagonizes the activity of the tuberous sclerosis complex 1/2 (TSC1/TSC2); TSC1/2 in turn inactivates RAS homologue enriched in brain (RHEB). The inactivation of RHEB disinhibits, i.e. activates, target of rapamycin (TOR). TOR then activates the effector gene *S6 kinase* (*S6K*) and inhibits the negative regulator 4E-BP. The phenotypic effects of naturally occurring alleles of the genes in the IIS/TOR pathway remain poorly understood, but clinal polymorphisms in *InR* (Paaby et al. 2010; Paaby et al. 2014) and *foxo* (this study) have pleiotropic effects on life history in *Drosophila*.

Since many fitness-related traits affected by IIS/TOR also exhibit phenotypic clines, it is tempting to hypothesize that natural variation in this pathway contributes to life-history clines, especially with regard to body size (de Jong and Bochdanovits 2003); yet, the evolutionary significance of natural variants in this pathway is poorly understood. An exception is an indel polymorphism in the *D. melanogaster InR* gene, which varies clinally along both the North American and Australian east coasts and which has multifarious life-history effects (Paaby et al. 2010, 2014). Consistent with the idea that IIS polymorphisms contribute to adaptation, natural variation in adult reproductive dormancy in *D. melanogaster* has been connected to the *Pi3K* gene (Williams et al. 2006), and work in *Caenorhabditis remanei* has identified a global selective sweep in the *Caenorhabditis* homolog of *Pi3K*, *age-1* (Jovelin et al. 2014). Multiple lines of evidence also indicate that insulin-like growth factor-1 (IGF-1) signaling mediates physiological life-history variation in vertebrate populations (Dantzer and Swanson 2011; Swanson and Dantzer 2014). Together, these findings suggest that allelic variation in IIS/TOR might profoundly affect life-history adaptation, but experimental evidence remains scarce (Flatt et al. 2013; Flatt and Partridge 2018).

Here we provide a comprehensive examination of the life-history effects of a clinally varying polymorphism in the forkhead box-O transcription factor gene *foxo* of *D. melanogaster* (Fig. 1), a major regulator of IIS that is homologous to *C. elegans daf-16* and mammalian *FOXO3A*. Molecular studies – mainly in the fly and nematode – have shown that FOXO plays a key role in regulating growth, lifespan and resistance to starvation and oxidative stress (Jünger et al. 2003; Puig et al. 2003; Libina et al. 2003; Murphy et al. 2003; Kramer et al. 2003; Kramer et al. 2008; Hwangbo et al. 2004; Puig and Tijan 2005; Fielenbach and Antebi 2008; Mattila et al. 2009; Slack et al. 2011). Moreover, genetic association studies in humans have linked polymorphisms in *FOXO3A* to longevity in centenarians (Flachsbart et al. 2009; Willcox et al. 2008). Natural *foxo* variants thus represent promising candidates for mediating life-history variation in natural populations.

From our previous population genomic data based on three population along the North American cline (Fabian et al. 2012) we identified two strongly clinally varying alternative *foxo* alleles, as defined by 2 focal SNPs, whose frequencies change across latitude by ~60% between Florida and Maine. Here we characterize the effects of these clinal *foxo* genotypes on several fitness-related traits (egg-to-adult survival, proxies of size, starvation resistance, fat content) by measuring phenotypes on replicate populations of the two alternative alleles under different environmental assay conditions in the laboratory. Since temperature gradients are thought to underpin – at least partly – latitudinal clines (e.g., de Jong and Bochdanovits 2003; Kapun et al. 2016b; and references therein), and because both diet and temperature modulate IIS (e.g., Britton et al. 2002; Kramer et al. 2003; Puig and Tijan 2005; Giannakou et al. 2008; Teleman 2010; Puig and Mattila 2011; Li and Gong 2015; Zhang et al. 2015), we phenotyped replicated population cage cultures of the alternative alleles at two temperatures (18°C, 25°C) and on two commonly used diets that differ mainly in their sugar source (sucrose vs. molasses) and content.

Measuring reaction norms to assess phenotypic plasticity and genotype-by-environment interactions (G × E) for this variant is interesting because still relatively little is known about whether and how clinality and plasticity interact (James et al. 1997; de Jong and Bochdanovits 2003; Hoffmann et al. 2005; Levine et al. 2011; Overgaard et al. 2011; Chen et al. 2012; Cooper et al. 2012; Zhao et al. 2015; Clemson et al. 2016; Mathur and Schmidt 2017; van Heerwaarden and Sgrò 2017). For example, *D. melanogaster* feeds and breeds on various kinds of rotting fruit, with the protein:carbohydrate (P:C) ratios exhibiting spatiotemporal variation (Lachaise et al. 1988; Hoffmann and McKechnie 1991; Markow et al. 1999; Keller 2007), but how dietary plasticity affects traits in a clinal context remains largely unclear. Similarly, the interplay between thermal plasticity and adaptation is incompletely understood (e.g., de Jong and Bochdanovits 2003; Overgaard et al. 2011; Mathur and Schmidt 2017; van Heerwaarden and Sgrò 2017).

We give predictions for the expected phenotypic effects of the *foxo* variant in terms of clinality, plasticity and the physiology of IIS in the Methods section. Our results show that the *foxo* polymorphism affects multiple fitness components according to these predictions and suggest that it contributes to clinal life-history adaptation; we also find that the alternative alleles respond plastically to temperature and diet but with little evidence for G x E.

## Methods

### IDENTIFICATION AND ISOLATION OF THE *FOXO* POLYMORPHISM

We identified two strongly clinal SNPs in *foxo* in the genomic data of Fabian et al. (2012) by using an *F*_ST_ outlier approach: an A/G polymorphism at position *3R*: 9892517 (position in the *D. melanogaster* reference genome v.5.0; *F*_ST_ = 0.48 between Florida and Maine) and a T/G polymorphism at position *3R*: 9894559 (*F*_ST_ = 0.42 between Florida and Maine) (Fig. S1A, Supporting Information; see Fabian et al. 2012 for details of outlier detection). The A/G polymorphism is a synonymous coding SNP, predicted to be located in the PEST region of the FOXO protein, which serves as a protein degradation signal (analysis with ExPASy [Artimo et al. 2012]; Fig. S2, Supporting Information). The T/G SNP is located in the first intron of *foxo*, with no biological function attributed to this position (Attrill et al. 2016).

While our initial identification of these SNPs was based on only three populations (Florida, Pennsylvania, Maine; Fabian et al. 2012), both SNPs are also strongly clinal in a more comprehensive dataset from 10 populations along the cline (Betancourt et al. 2018), collected by the *Drosophila* Real Time Evolution Consortium (DrosRTEC; Bergland et al. 2014; Kapun et al. 2016b; Machado et al. 2018). The frequency of the high-latitude [HL] allele (A, T) for this 2-SNP variant ranges from ~10% in Florida to ~70% in Maine; conversely, the alternative low-latitude [LL] allele (G,G) is prevalent in Florida but at low frequency in Maine (Fig. S1A, Supporting Information). Because the two *foxo* SNPs are located relatively closely to each other (~2 kb apart; Fig. S1A, Supporting Information), we decided to study them experimentally in combination, as alternative 2-SNP alleles. Indeed, as shown in Fig. S1B (Supporting Information), the two focal *foxo* SNPs are in perfect linkage disequilibrium (LD; *r*^2^ = 1), without any significant LD in-between the two sites. Although these SNPs do not fall outside the empirical neutral distribution (based on 20,000 SNPs in short introns), they do fall in the tails of the distribution (Betancourt et al. 2018); further work will thus be required to assess whether the allele frequency observations are consistent with neutral demographic history or if a model of spatially varying or balancing selection needs to be invoked.

To isolate the two alternative *foxo* alleles for experiments we used whole-genome sequenced inbred lines from the *Drosophila* Genetic Reference Panel (DGRP; Mackay et al. 2012) to reconstitute outbred populations either fixed for the LL (G,G) and the HL (A,T) alleles. This ‘reconstituted or recombinant outbred population’ (ROP) or ‘Mendelian randomization’ approach produces populations that are consistently and completely fixed for the two alternative allelic states to be compared, with the rest of the genetic background being randomized (see Behrman et al. 2018 and Lafuente et al. 2018 for examples using this method). For each allele we used two independent sets of DGRP lines (sets A and B for HL; sets C and D for LL; each set consisting of 20 distinct lines) and two replicate population cages per set, giving a total of 8 population cages (Fig. S3, Table S1, Supporting Information). ROP cages were established from the DGRP lines by Betancourt et al. (2018); F2 flies were transferred to Lausanne for establishing cages at our laboratory.

**Table 1.** Summary of ANOVA results for viability; femur length; wing area:thorax length ratio; female starvation resistance (also cf. Table S5). White and grey cells show results for females and males, respectively. \**p* < 0.05; \*\**p* < 0.01; \*\*\**p* < 0.001.

| Factor | Proportion Viability | Femur Length | Wing Area: Thorax Length Ratio | Starvation Resistance |
| --- | --- | --- | --- | --- |
| Allele | $F_{1,32}=20.65^{***}$ | $F_{1,32}=16.66^{***}$ | $F_{1,4}=46.64^{***}$ | $F_{1,32}=23.86^{***}$ |
| | | $F_{1,32}=0.16$ | $F_{1,4}=82.17^{***}$ | |
| Temperature | $F_{1,114}=3.24$ | $F_{1,1923}=1617.80^{***}$ | $F_{1,18}=477.45^{***}$ | $F_{1,1547}=732.08^{***}$ |
| | | $F_{1,1923}=443.60^{***}$ | $F_{1,18}=1366.87^{***}$ | |
| Diet | $F_{1,114}=8.43^{**}$ | $F_{1,1923}=144.72^{***}$ | $F_{1,18}=50.35^{***}$ | $F_{1,1547}=129.99^{***}$ |
| | | $F_{1,1923}=68.24^{***}$ | $F_{1,18}=127.77^{***}$ | |
| Allele x Temperature | $F_{1,114}=2.25$ | $F_{1,1923}=0.36$ | $F_{1,18}=0.14$ | $F_{1,1547}=3.43$ |
| | | $F_{1,1923}=1.40$ | $F_{1,18}=0.32$ | |
| Temperature x Diet | $F_{1,114}=1.85$ | $F_{1,1923}=13.26^{***}$ | $F_{1,18}=16.64^{***}$ | $F_{1,1547}=14.81^{***}$ |
| | | $F_{1,1923}=4.65$ | $F_{1,18}=56.36^{***}$ | |
| Allele x Diet | $F_{1,114}=1.71$ | $F_{1,1923}=3.28$ | $F_{1,18}=0.21$ | $F_{1,1547}=16.22^{***}$ |
| | | $F_{1,1923}=4.04^{*}$ | $F_{1,18}=2.53$ | |
| Allele x Temperature x Diet | $F_{1,114}=0.39$ | $F_{1,1923}=6.41^{*}$ | $F_{1,18}=0$ | $F_{1,1547}=1.63$ |
| | | $F_{1,1923}=0.95$ | $F_{1,18}=8.34^{**}$ | |
| Set(Allele) | $F_{2,32}=2.50$ | $F_{2,32}=5.89^{**}$ | $F_{2,4}=6.86^{**}$ | $F_{2,32}=45.24^{***}$ |
| | | $F_{2,32}=0.75$ | $F_{2,4}=3.80^{*}$ | |
| Cage(Set, Allele) | $F_{4,32}=61.25^{***}$ | $F_{4,32}=37.43^{***}$ | NA | $F_{4,32}=11.17^{***}$ |
| | | $F_{4,32}=415.66^{***}$ | NA | |

Genomic analysis of the DGRP lines used to set up the ROP cages showed that sets A and B versus sets C and D were completely fixed (*F*_ST_ = 1) for the HL and LL alleles, respectively; this also showed that, although there exist other SNPs that are strongly differentiated (*F*_ST_ >0.5) between the HL and LL populations, the majority of them are different between the independently replicated sets (blocks) of DGRP lines used to make the HL vs. LL contrast (Betancourt et al. 2018). Such SNPs, which are specific to a given set of lines, do not make a consistent contribution to the overall HL vs. LL contrast.

The most parsimonious interpretation of our results is therefore that the effects reported below are caused by the two *foxo* SNPs which we have studied. However, we cannot completely rule out that other (causative) sites are potentially in long-range LD with our focal SNPs (see Fig. S1B, Supporting Information). A conservative interpretation of our results is thus to view the two focal *foxo* SNPs as representing ‘tags’ or markers for functionally significant variants segregating at the *foxo* locus that are in LD with the causative site(s), similar to those used in genome-wide association studies (GWAS; e.g., Wang et al. 2010).

### POPULATION CAGES

Population cages were maintained at 25°C, 12:12 h light:dark, 60% relative air humidity and controlled larval density. Larval density was kept constant via egg collections (200-300 eggs per bottle [6 oz. = 177 mL]; 10 bottles per cage), with eclosing adults being released into cages (17.5 x 17.5 x 17.5 cm; BugDorm®) at a density of ~2000-2500 adults per cage. Prior to the phenotypic assays population cages were kept for 10 generations to allow for free recombination among lines within each cage and allelic state and to homogenize (randomize) differences in genomic background between the two allelic states to be compared. Before setting up assays, we kept cages for 2 generations under common garden conditions (room temperature: ~22°C, ~10:14 h light:dark, ~50% humidity). Thus, phenotypes were measured after a total of 12 generations of recombination.

### PHENOTYPE ASSAYS

Assays reported here were performed in our laboratory in Lausanne; independent assays were performed under constant environmental conditions by Betancourt et al. (2018), thus allowing us to assess the reproducibility of our results and to identify potential differences in assay conditions between laboratories (cf. Ackermann et al. 2001).

In generation 13 (see above) we assayed flies for viability, size, starvation resistance and lipid content. Phenotypes were assayed under four environmental conditions, using a fully factorial 2-way design: 2 rearing temperatures (18°C, 25°C) by 2 commonly used diets that differ mainly in their sugar source (sucrose [cornmeal-agar-yeast-sucrose] vs. molasses [cornmeal-agar-yeast-molasses] diet and their protein:carbohydrate ratio (P:C ~1:3.6 vs. ~1:12.3, respectively; see Table S2, Supporting Information, for details of nutrient content and media recipes). To initiate assays we collected ~6400 eggs from each cage, distributed them across 32 bottles (each with 200 eggs; 25 mL medium), and allocated 8 bottles to each of the 4 conditions (8 bottles × 8 cages × 4 conditions = 256 bottles). For all assays (except viability; see below), we collected eclosed adults in 48-h cohorts, allowed them to mate for 4 days under their respective thermal and dietary conditions, sexed them under light CO_2_ anesthesia 4-6 days post-eclosion, and transferred them to fresh vials 24 h prior to assays. Flies used for size assays were stored at −20°C until measurement.

**Table 2.** ANOVA results for female fat loss upon starvation. \**p* < 0.05; \*\**p* < 0.01; \*\*\**p* < 0.001. The fixed factor ‘Treatment’ has two levels: fed vs. starved; interactions involving the factors ‘Allele’ and ‘Treatment’ test for allelic differences in fat catabolism.

| Factor | Fat content |  |
| --- | --- | --- |
|  | 18°C | 25°C |
| Allele | $F_{1,32}=0.02$ | $F_{1,32}=1.90$ |
| Diet | $F_{1,301}=70.97^{***}$ | $F_{1,300}=310.82^{***}$ |
| Treatment | $F_{1,301}=223.48^{***}$ | $F_{1,300}=130.68^{***}$ |
| Allele x Diet | $F_{1,301}=20.58^{***}$ | $F_{1,300}=6.93^{**}$ |
| Diet x Treatment | $F_{1,301}=25.46^{***}$ | $F_{1,300}=21.31^{***}$ |
| Allele x Treatment | $F_{1,301}=7.01^{**}$ | $F_{1,300}=1.24$ |
| Allele x Diet x Treatment | $F_{1,301}=0$ | $F_{1,300}=7.03^{**}$ |
| Set(Allele) | $F_{2,32}=13.11^{***}$ | $F_{2,32}=4.24^*$ |
| Cage(Set, Allele) | $F_{4,32}=9.46^{***}$ | $F_{4,32}=1.44$ |

Viability (egg-to-adult survival) was calculated as the proportion of adult flies successfully developing from eggs by collecting 600 eggs per cage and placing them into vials containing 8 mL of medium, with 30 eggs per vial (5 vials × 8 cages × 4 conditions = 160 vials).

Body size was examined by measuring three proxies: wing area, thorax length and femur length (*N* = 26-30 wings, 9-15 thoraces, and 19-21 femurs per cage, treatment, and sex). Right wings and femurs were mounted on slides with CC/Mount^™^ tissue mounting medium (Sigma Aldrich) and slides sealed with cover slips. Thorax length was defined as the lateral distance between the upper tip of the thorax and the end of the scutellar plate (*N* = 10-15 individuals per cage, treatment, and sex). Images for morphometric measurements were taken with a digital camera (Leica DFC 290) attached to a stereo dissecting microscope (Leica MZ 125; Leica Microsystems GmbH, Wetzlar, Germany). We used ImageJ software (v.1.47) to measure femur and thorax length (mm) and to define landmarks for calculating wing area (mm^2^). To measure wing area we defined 12 landmarks located at various vein intersections along the wing; the total area encompassed by these landmarks was estimated using a custom-made Python script (available upon request). In brief, we split the polygon defined by the landmarks up into triangles and summed across their areas (Fig. S4, Supporting Information). Thorax and femur (but not wing area) measurements were repeated three times per individual (see below for estimates of ‘repeatability’). From these data, we calculated the ratio of wing area:thorax length, which is inversely related to ‘wing loading’ (Azevedo et al. 1998; Gilchrist et al. 2000); reduced wing loading (i.e., increased wing dimensions relative to body size) can improve flight performance at low temperature (Frazier et al. 2008).

To measure starvation resistance (i.e., survival upon starvation) we placed flies into vials containing 0.5% agar/water medium and scored the duration of survival (h) upon starvation every 6 h until all flies had died (*N* = 5 vials × 10 flies per vial × 2 sexes × 8 cages × 4 conditions = 320 vials or 3200 flies).

Since there is typically a positive correlation between starvation resistance and lipid content (Hoffmann and Harshman 1999), we also determined whole-body triacylglyceride (TAG) content (in *µ*g per fly) using a serum triglyceride determination kit (Sigma Aldrich; Tennessen et al. 2014). For each cage and treatment, triglyceride content was estimated from 5-7-day-old females, either kept under fed or starved (24 h) conditions, by preparing 10 replicate homogenates, each made from 2 flies (8 cages × 4 conditions × 2 treatments × 10 replicates = 640 homogenates). To estimate fat loss upon starvation we calculated the difference between fat content under fed versus starved conditions, using treatment (fed vs. starved) means from each population cage (mean fat loss per fly, in *µ*g).

### QRT-PCR ANALYSIS OF INSULIN SIGNALING STATE

A well-established transcriptional read-out of FOXO signaling is the insulin-like receptor InR: under conditions of high insulin (e.g., after a meal), InR synthesis is repressed by a feedback mechanism controlled by FOXO; conversely, under conditions of low insulin, activation of FOXO leads to upregulation of *InR* mRNA (Puig et al. 2003; Puig and Tjian 2005). To test whether the *foxo* alleles differ in IIS state we performed qRT-PCR, measuring *InR* mRNA abundance. For each cage and treatment, we extracted total RNA from 5-7-day-old snap-frozen females in triplicate, with each replicate prepared from 5 flies. RNA was extracted with the RNeasy kit (Qiagen) and reverse transcribed with the GoScript Reverse Transcription System (Promega). From each triplicate biological sample we prepared 3 technical replicates (8 cages × 4 conditions × 3 biological replicates × 3 technical replicates = 288 samples). Relative transcript abundance was normalized by using *Actin5C* as an endogenous control (Ponton et al. 2011). qRT-PCR was carried out using a QuantStudio 6 Flex Real-Time PCR System (Applied Biosystems) and SYBR Green GoTaq qPCR Master Mix (Promega). Thermal cycling was conducted at 95°C for 2 min, followed by 42 cycles of amplification at 95°C for 15 s and 60°C for 1 min, and using the following melting curve: 95°C for 15 s, 60°C for 1 min, and 95°C for 15 s. Quantification of relative abundance for each sample was based on the Δ*CT* method. We used the following primer sequences (Casas-Tinto et al. 2007; Ponton et al. 2011): *Actin5C forward*, 5’-GCGTCGGTCAATTCAATCTT-3’; *Actin5C reverse*, 5’-AAGCTGCAACCTCTTCGTCA-3’; *InR forward*, 5’-CACAAGCTGGAAAGAAAGTGC-3’; *InR reverse*, 5’-CAAACACGTTTCGATAATATTTTTCT-3’.

### STATISTICAL ANALYSIS

Analyses were performed with JMP (SAS, Raleigh, NC, USA; v.11.1.1). Data were analyzed with analysis of variance (ANOVA), testing the fixed effects of allele (*A*; HL vs. LL), temperature (*T*; 18°C vs. 25°C), diet (*D*; sucrose vs. molasses), set (*S*; independent blocks of DGRP lines) nested within *A*, replicate cage (*C*) nested within the combination of *A* and *S*, and all 2- and 3-way interactions: *y = A + T + D + A × T + A × D + T × D + A × T × D + S(A) + C(A,S)*, where *y* denotes the response variable (trait). For simplicity, the sexes were analyzed separately (i.e., to reduce the number of higher-order interactions).

For starvation resistance we measured age at death from multiple individuals per replicate vial; we thus estimated and accounted for the random effect of vial (*V*), nested within the combination of *A*, *S* and *C*, using restricted maximum likelihood (REML) (see Supporting Information for these estimates).

Viability (proportion) data were arcsine square-root transformed prior to analysis. ANOVA on thorax and femur length data was performed using means across 3 measures per individual. From the repeat measurements of these traits on the same individuals, we estimated the ‘repeatability’ of our measurements (i.e., the intraclass correlation; see Whitlock and Schluter 2009) by performing random-effects ANOVAs with REML. Overall, repeatibility was very high for femur length (~91.9% for females; 94.4% for males) but less so for thorax length (~29.9% for females; 36.6% for males) (details not shown). Because wings and thoraces were measured on separate individuals, analysis of wing:thorax ratio was performed on population (cage) means. For fat content, we included the fixed effect of starvation treatment (*Tr*; fed vs. starved); interactions involving *A* and *Tr* (i.e., *A × Tr*; *A × D × Tr*) test for allelic differences in fat loss upon starvation. For simplicity, this analysis was performed separately for the two rearing temperatures.

To estimate the magnitude of the allelic effects of the *foxo* polymorphism upon the assayed fitness components we calculated Cohen’s *d* (Table S3, Supporting Information), a standardized measure of effect size (i.e., a signal to noise ratio, defined as the difference between two means divided by their pooled standard deviation) (Cohen 1988; Sawilowsky 2009). Low values of Cohen’s *d* (e.g., 0.01) are commonly interpreted as representing very small effect sizes, whereas effect sizes >0.8 are interpreted as being qualitatively large to very large (Sawilowsky 2009).

We also estimated the relative contribution of the *foxo* polymorphism assayed in our laboratory to the overall phenotypic cline for wing area, recently measured on flies from 6 populations along the North American east coast (Betancourt et al. 2018; using flies assayed on molasses diet at 25°C). We calculated the proportional contribution of the polymorphism to the overall cline as follows: Δ*_foxo_* × Δ_frequency_ */* Δ_cline_, where Δ*_foxo_* is the difference in mean wing area between the HL and LL alleles, Δ_frequency_ is the allele frequency gradient for the polymorphism between cline ends (Maine vs. Florida, ~60%) and Δ_cline_ is the difference in mean wing area between cline ends.

### PREDICTIONS

Here we make some qualitative predictions for the expected behavior of the *foxo* polymorphism with regard to (1) clinal phenotypic effects, (2) patterns of trait covariation determined by IIS, and (3) plasticity, G × E, and local adaptation. We compare our results to these predictions in the Results section below.

(1) Latitudinal clinality. Traits which have been found to covary positively with latitude include, for example, faster development, lower egg-to-adult survival (viability), increased body size, reduced wing loading, reduced fecundity, prolonged lifespan, and increased resistance to starvation, cold and heat stress (e.g., Coyne and Beecham 1987; Azevedo et al. 1998; Bochdanovits and de Jong 2003a; de Jong and Bochdanovits 2003; Schmidt et al. 2005a, 2005b; Folguera et al. 2008; Schmidt and Paaby 2008; Bhan et al. 2014; Mathur and Schmidt 2017; Durmaz et al. 2018). For some traits clinal patterns have been observed in a parallel fashion on multiple continents, but there can also be major differences among continents (e.g., see discussion in Fabian et al. 2015); for example, contrasting predictions have been made for viability (van ‘t Land et al. 1999), starvation resistance (Karan et al. 1998, Robinson et al. 2002; Hoffmann et al. 2005; Goenaga et al. 2013) and heat tolerance (Hoffmann et al. 2002; Sgrò et al. 2010).

In general, we would expect that the effects of the high- and low-latitude *foxo* alleles agree with the overall phenotypic patterns across latitude, especially for those traits that have previously been examined along the North American cline (e.g., Coyne and Beecham 1987; Schmidt and Paaby 2008; Paaby et al. 2014; Kapun et al. 2016a; Mathur and Schmidt 2017; Durmaz et al. 2018).

(2) IIS. Traits that are associated with reduced IIS include reduced body size, increased lifespan, resistance to starvation and cold, increased fat content, reduced fecundity, and activation of FOXO (Tatar et al. 2001, 2003; Oldham and Hafen 2003; Broughton et al. 2005; Teleman 2010). For example, loss-of-function (LOF) mutants of *foxo* exhibit (depending on the allele) prolonged development, reduced weight, smaller wing size, reduced fecundity, shortened lifespan, and reduced survival upon oxidative and starvation stress (Jünger et al. 2003; Kramer et al. 2003, 2008; Hwangbo et al. 2004; Giannakou et al. 2004; Giannakou et al. 2008; Slack et al. 2011); the effects of IIS (or of *foxo*) on viability are, however, not well understood. Conversely, overexpression of *foxo* has opposite effects on most of these traits (e.g., increased lifespan), yet – like LOF alleles – causes decreased size (Kramer et al. 2003; Puig et al. 2003; Hwangbo et al. 2004; Kramer et al. 2008; Tang et al. 2011).

We predict that the naturally occurring *foxo* alleles tested here differ consistently along this IIS/*foxo* axis of trait covariation. Notably, traits observed in flies from high-versus low-latitude populations in North America resemble those of flies with low versus high IIS, respectively (e.g., de Jong and Bochdanovits 2003; Flatt et al. 2013; Paaby et al. 2014): lower fecundity, improved stress resistance, and longer lifespan observed in high-latitude flies are traits that tend to be co-expressed in IIS mutants; however, flies from high-latitude populations are larger than low-latitude flies, yet reduced IIS causes smaller size.

(3) Plasticity, G × E, and local adaptation. With regard to thermal effects, we would expect flies raised at lower temperature to exhibit prolonged development, reduced viability, larger size, reduced wing loading, lower fecundity, increased lifespan, and improved starvation resistance (David et al. 1994; Partridge et al. 1994a, 1994b; James and Partridge 1995; Bochdanovits and de Jong 2003b; Trotta et al. 2006; Folguera et al. 2008; Klepsatel et al. 2013, 2014; Mathur and Schmidt 2017; cf. Hoffmann et al. 2005 for a contrasting prediction for starvation survival).

With respect to dietary effects, higher P:C ratios, for instance, might be expected to cause increased viability, larger size but reduced starvation resistance (Lee and Jang 2014; Lihoreau et al. 2016; Reis 2016). In terms of G × E, genotypes from temperate, seasonal high-latitude habitats might be more plastic than those from low-latitude habitats (Overgaard et al. 2011; Klepsatel et al. 2013); if so, patterns of differential plasticity between high- and low-latitude alleles might be consistent with patterns of local adaptation (Mathur and Schmidt 2017).

## Results

The clinal *foxo* polymorphism examined here (or causative SNPs in LD with it; see caveat in the Methods section) impacted all fitness components assayed (Table 1; Tables S3 and S4, Supporting Information), including significant effects on egg-to-adult survival (viability) (qualitatively moderate to large effects, as measured by Cohen’s *d*), femur length (very small to medium), wing area (medium), thorax length (very small to very large), starvation resistance (very small to medium), and lipid content (very small to large effects).

### ALLELIC VARIATION AT *FOXO* AFFECTS VIABILITY

The *foxo* polymorphism significantly affected viability, with the LL allele exhibiting higher egg-to-adult survival than the HL allele (Fig. 2; Table 1), consistent with observations suggesting that viability might be higher at low latitudes (Folguera et al. 2008; but see van ‘t Land et al. 1999). Diet – but not temperature – also had an effect, with viability being higher on sucrose than on molasses diet (Fig. 2; Table 1). We did not find any evidence for G × E interactions affecting this trait.

**Figure 2.**
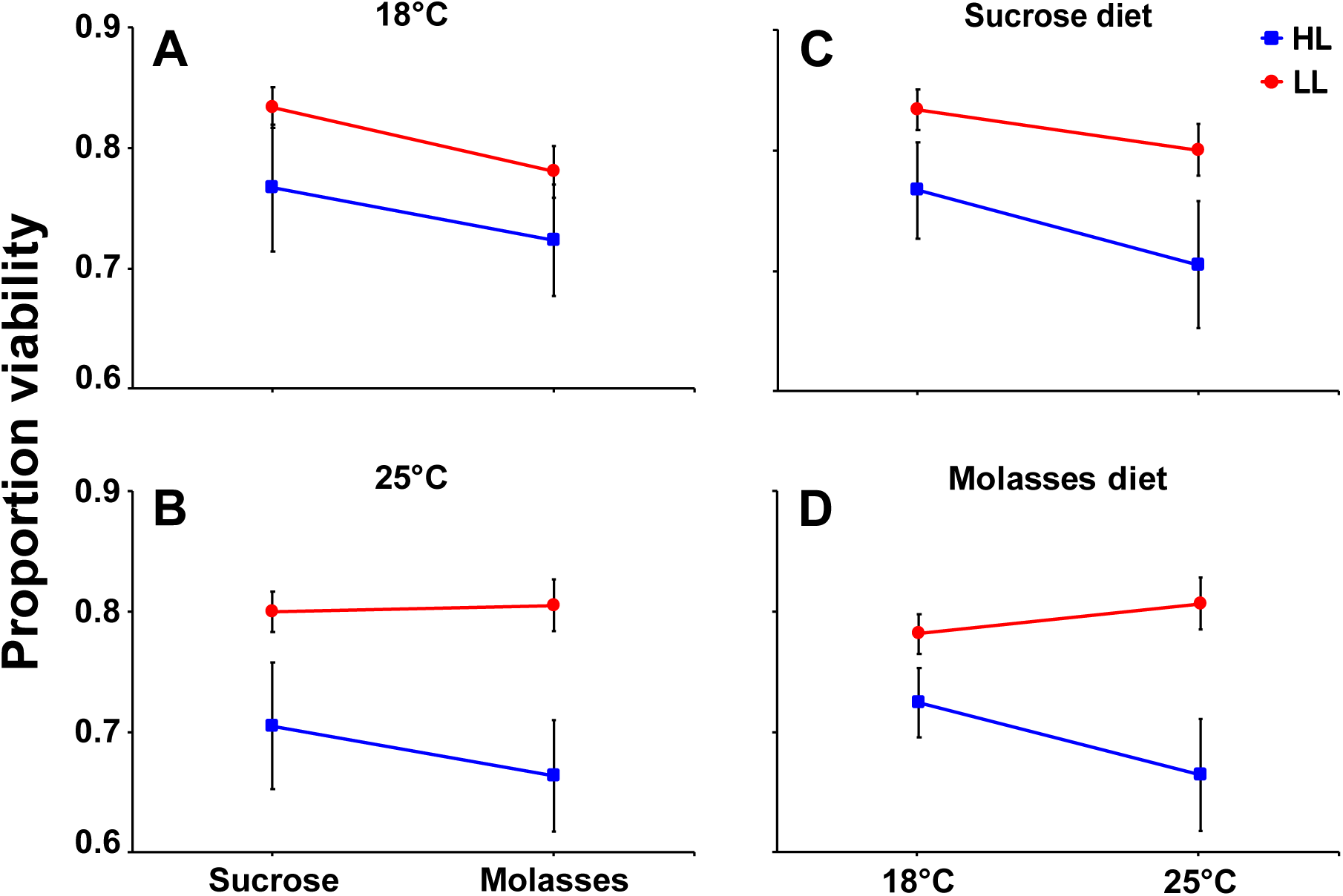
Viability (egg-to-adult survival). Effects of the clinal *foxo* variant on the proportion viability (egg-to-adult survival). (A) Dietary reaction norms at 18°C. (B) Dietary reaction norms at 25°C. (C) Thermal reaction norms measured on sucrose diet. (D) Thermal reaction norms measured on molasses diet. Data in (A, B) are the same as those shown in (C, D). Shown are means and standard errors. Red lines: low-latitude (LL) allele, blue lines: high-latitude (HL) allele.

### CLINAL *FOXO* ALLELES DIFFER IN BODY SIZE

Since both latitude and IIS affect size (de Jong and Bochdanovits 2003), we next examined three proxies of body size (wing area, thorax and femur length). The HL allele conferred larger femur length (Fig. 3; Table 1; in females but not males), wing area (Fig. S5; Table S4, Supporting Information), and wing:thorax ratio than the LL allele (Fig. 4; Table 1; for thorax data see Fig. S6; Table S4, Supporting Information). These results are consistent with the positive size cline in North America (Coyne and Beecham 1987) and with reduced wing loading at high latitude (Azevedo et al. 1998; Bhan et al. 2014). Remarkably, with regard to wing area, we estimate that the *foxo* polymorphism makes a proportional contribution of ~14% to the total cline for wing area (females: Δ*_foxo_* × Δ_frequency_ / Δ_cline_ ≈ 0.017 × 0.6 / 0.074 ≈ 0.138; males: Δ*_foxo_* × Δ_frequency_ / Δ_cline_ ≈ 0.019 × 0.6 / 0.083 ≈ 0.137) – this represents a major contribution to the wing size cline along the North American east coast (Coyne and Beecham 1987; Betancourt et al. 2018).

**Figure 3.**
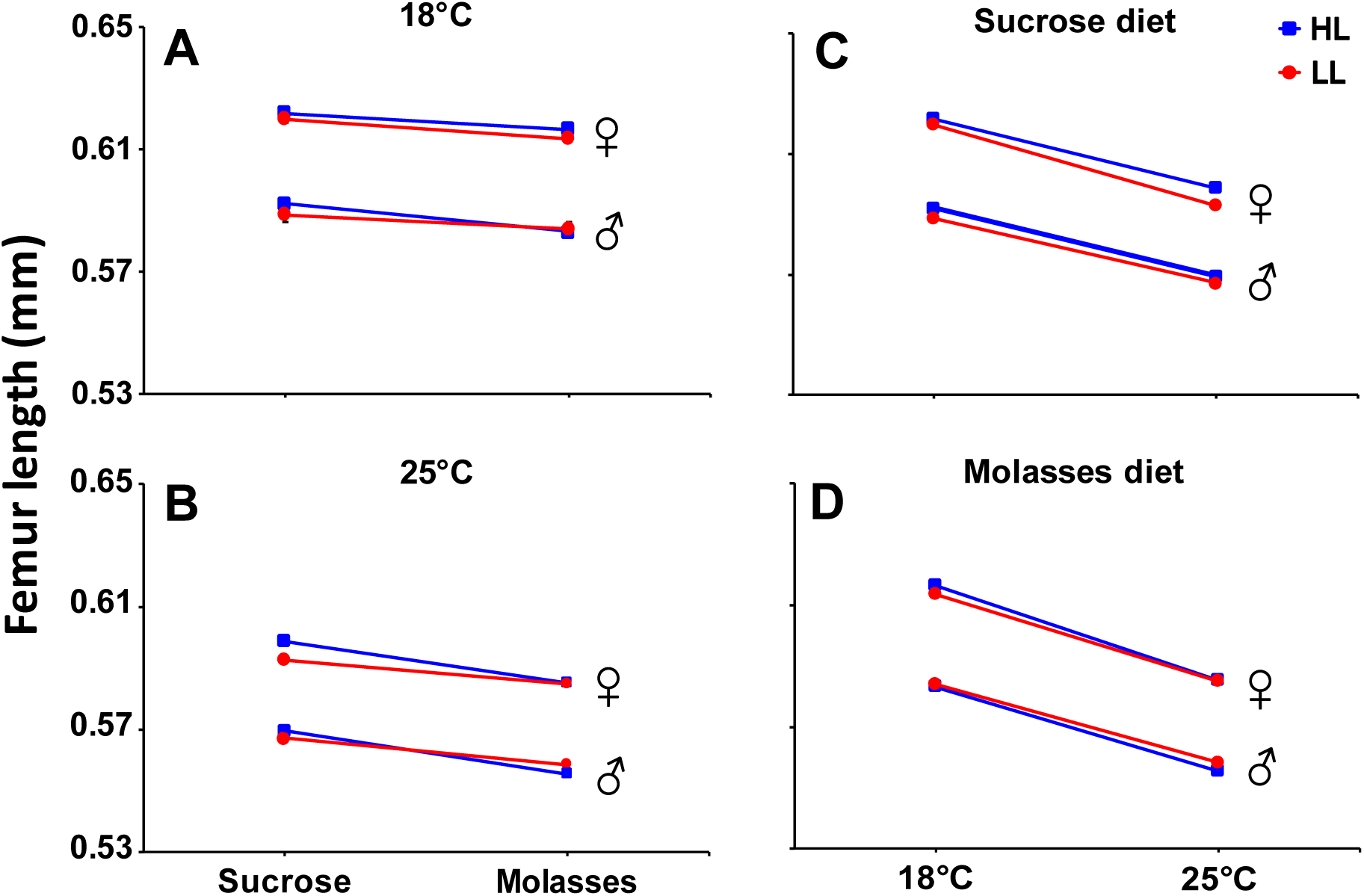
Femur length. Effects of the *foxo* polymorphism on femur length (mm) in females and males. (A) Dietary reaction norms at 18°C. (B) Dietary reaction norms at 25°C. (C) Thermal reaction norms measured on sucrose diet. (D) Thermal reaction norms measured on molasses diet. Data in (A, B) are the same as those shown in (C, D). Shown are means and standard errors. Red lines: low-latitude (LL) allele, blue lines: high-latitude (HL) allele.

**Figure 4.**
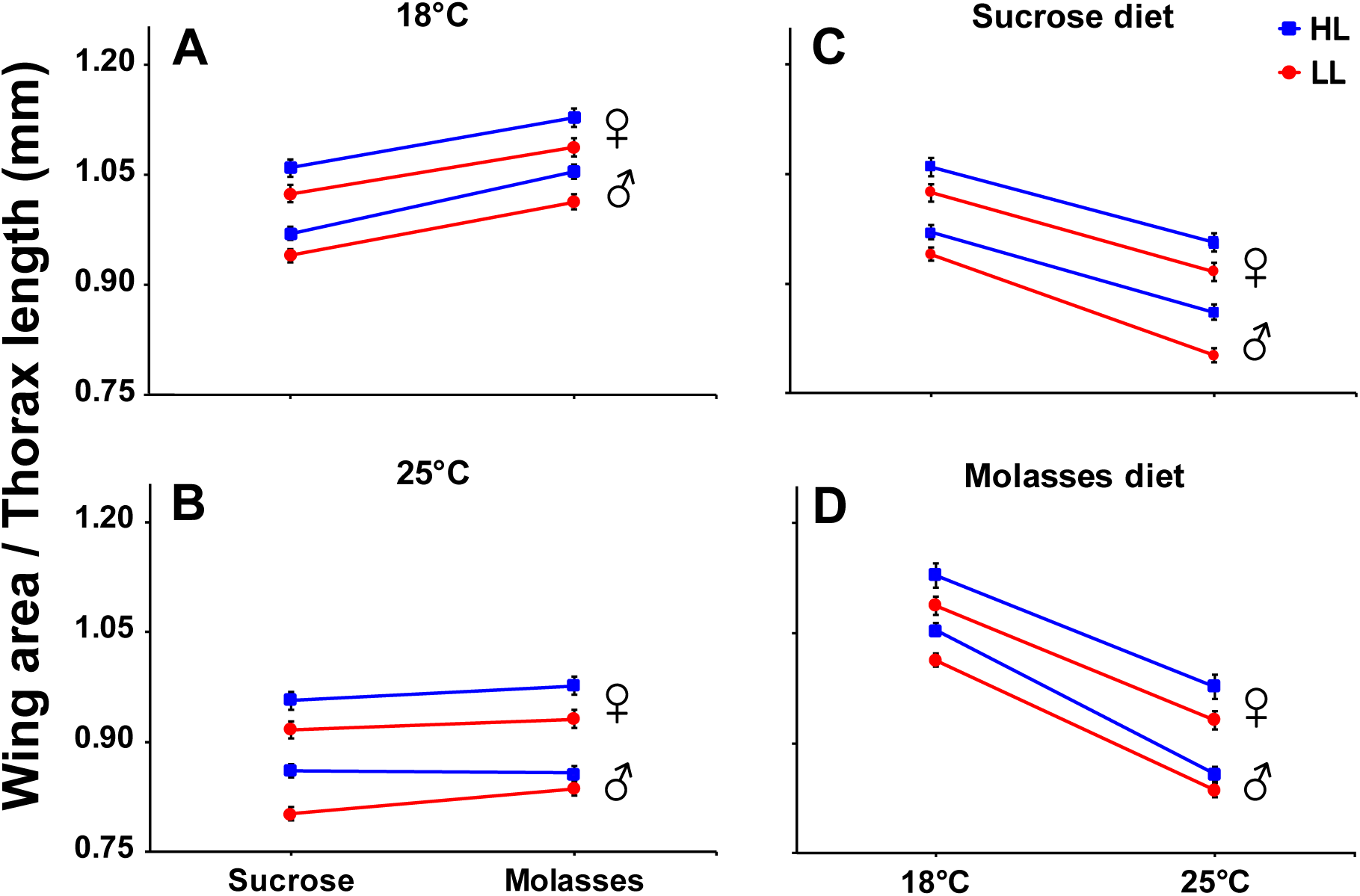
Wing:thorax ratio. Effects of the *foxo* variant on the ratio of wing area:thorax length (mm) in females and males. (A) Dietary reaction norms at 18°C. (B) Dietary reaction norms at 25°C. (C) Thermal reaction norms measured on sucrose diet. (D) Thermal reaction norms measured on molasses diet. Data in (A, B) are the same as those shown in (C, D). Shown are means and (propagated) standard errors. Red lines: low-latitude (LL) allele, blue lines: high-latitude (HL) allele.

For all size traits, females were larger than males (Fig. 3; Fig. 4; Table 1; Fig. S5; Fig. S6; Table S4, Supporting Information), as is typically observed. With regard to the plastic effects of temperature, femur length, thorax length and wing area were larger at 18°C than at 25°C (Fig. 3; Fig. S5, Fig. S6, Supporting Information; Table 1; Table S4, Supporting Information), as is expected based on previous work (David et al. 1994; Partridge et al. 1994a). In terms of dietary plasticity, femur and thorax length were larger on sucrose than on molasses diet (Fig. 3; Table 1; Fig. S6; Table S4, Supporting Information), perhaps in line with the observation that more carbohydrate-rich diets cause smaller size (Reis 2016); however, wing area and wing:thorax ratio were larger on molasses than on sucrose diet (Fig. S5; Table S4, Supporting Information; and Fig. 4; Table 1). Although we found a few G × E interactions for size traits (Fig. 4; Fig. 5; Table 1; Fig. S5; Fig. S6; Table S4, Supporting Information), the allelic reaction norms were overall remarkably parallel across environmental conditions.

**Figure 5.**
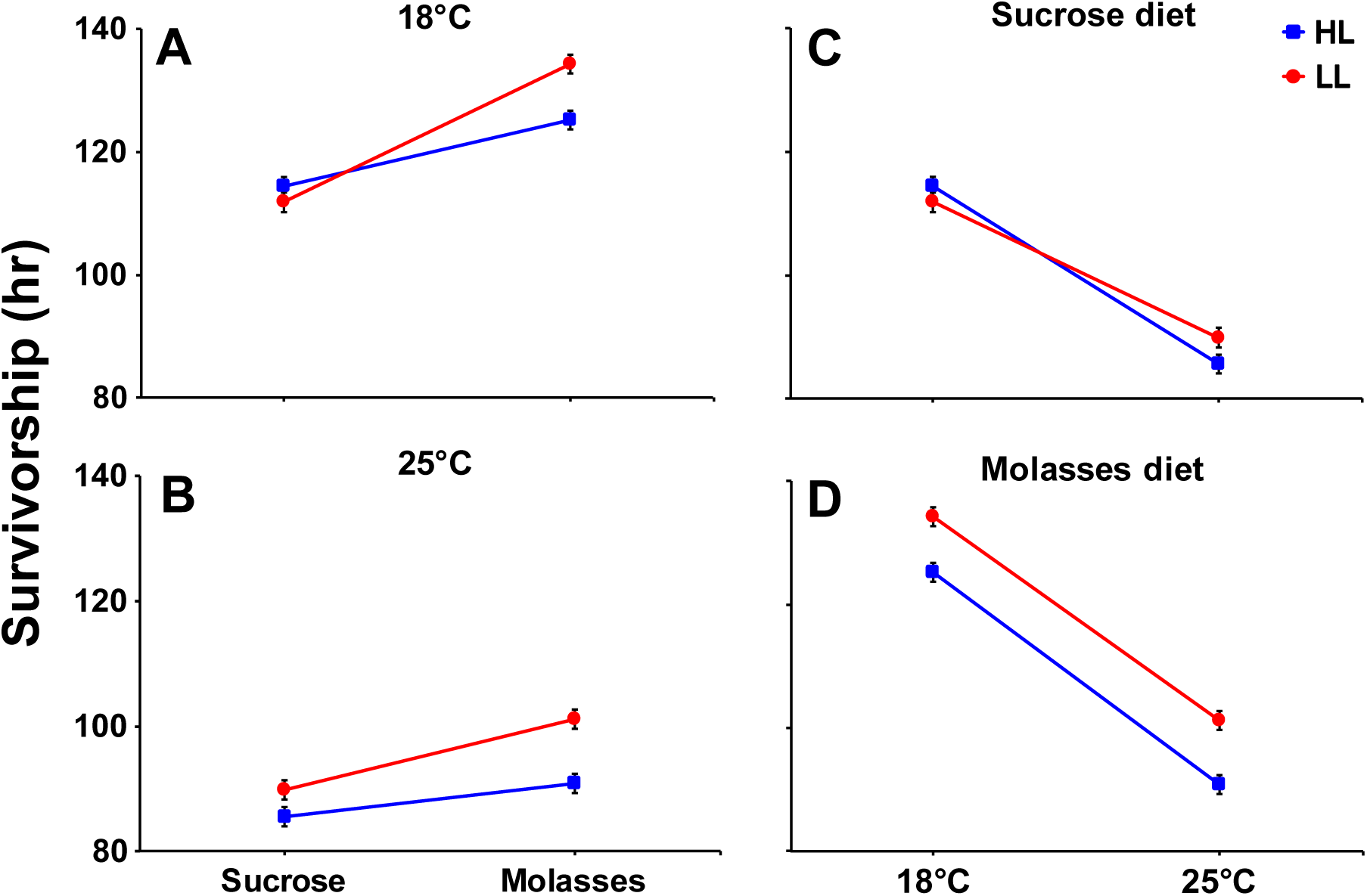
Starvation resistance. Effects of the clinal *foxo* polymorphism on the duration of survival (in hrs) upon starvation in females. (A) Dietary reaction norms at 18°C. (B) Dietary reaction norms at 25°C. (C) Thermal reaction norms measured on sucrose diet. (D) Thermal reaction norms measured on molasses diet. Data in (A, B) are the same as those shown in (C, D). Shown are means and standard errors. Red lines: low-latitude (LL) allele, blue lines: high-latitude (HL) allele.

### POLYMORPHISM AT *FOXO* IMPACTS STARVATION AND FAT CATABOLISM

The *foxo* alleles also differed in their effects on resistance to (survival of) starvation in females (Fig. 5; Table 1), as might be expected based on the observation that *foxo* mutants are more starvation sensitive than wildtype (Jünger et al. 2003; Kramer et al. 2003, 2008). However, contrary to clinal predictions (e.g., Schmidt and Paaby 2008; Mathur and Schmidt 2017), LL females were more resistant than HL females (Fig. 5; Table 1), suggesting a countergradient effect; in males, there were no allelic differences in resistance (Fig. S7; Table S4, Supporting Information; for estimates of the variance components of the random effect of vial see Table S5, Supporting Information). Overall females were more resistant than males (Fig. 5; Table 1; Fig. S7; Table S4, Supporting Information), consistent with some but not other studies (Goenaga et al. 2010; but see Matzkin et al. 2009). For both females and males, starvation resistance was higher at 18°C than at 25°C (Fig. 5; Table 1; Fig. S7; Table S4, Supporting Information), as previously reported (Mathur and Schmidt 2017). Flies raised on molasses diet were more resistant than those raised on sucrose diet (Fig. 5; Table 1; Fig. S7; Table S4, Supporting Information), potentially in support of the finding that lower P:C ratios favor higher resistance (Chippindale et al. 1993; Lee and Jang 2014). We also found evidence for an allele by diet interaction: allelic differences in resistance were more pronounced on molasses than sucrose diet (Fig. 5; Table 1; Fig. S7; Table S4, Supporting Information).

To further examine the physiological basis of starvation resistance we quantified how much fat female flies mobilize upon starvation (Fig 6; Table 2; males were not examined since they did not show allelic differences in resistance). Paralleling our result that LL females are more resistant than HL females, the amount of fat catabolized under starvation was greater in LL than in HL females, under almost all conditions (except for females raised on sucrose diet at 25°C; see Fig. 6 and Table 2: significant allele by diet by starvation treatment interaction at 25°C but not at 18°C). Fat loss upon starvation was greater for flies raised on molasses than on sucrose diet (Fig 6; Table 2), again matching the results for starvation resistance itself.

**Figure 6.**
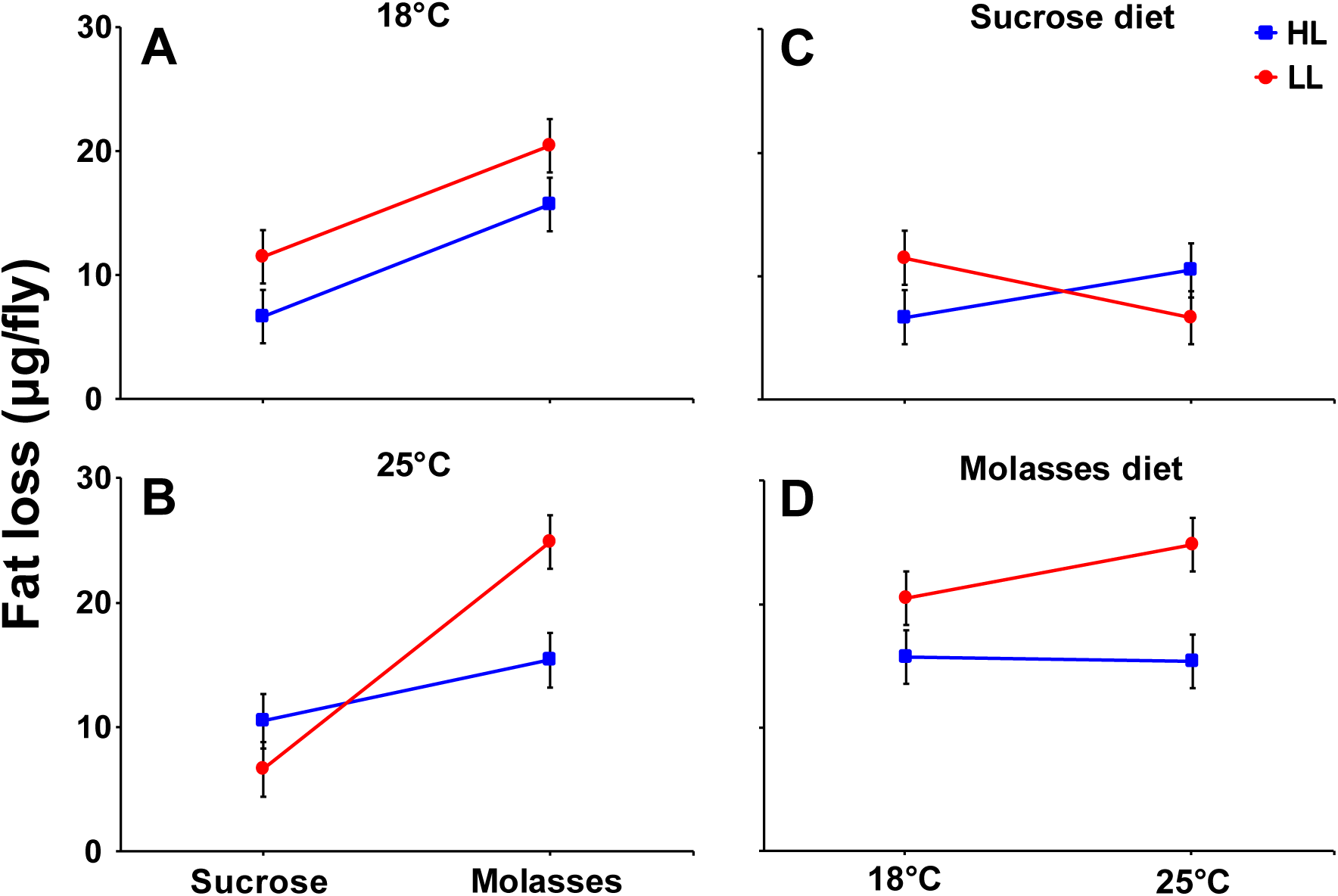
Fat loss upon starvation. Effects of the clinal *foxo* variant on female triglyceride loss upon starvation (*µg*/fly). (A) Dietary reaction norms at 18°C. (B) Dietary reaction norms at 25°C. (C) Thermal reaction norms measured on sucrose diet. (D) Thermal reaction norms measured on molasses diet. Data in (A, B) are the same as those shown in (C, D). Shown are means and (propagated) standard errors. Red lines: low-latitude (LL) allele, blue lines: high-latitude (HL) allele.

### *FOXO* ALLELES DIFFER IN TRANSCRIPTIONAL FEEDBACK CONTROL OF *InR*

From the above patterns we predicted that the LL allele would exhibit decreased IIS and increased FOXO activity: the LL allele has smaller size but higher starvation resistance, i.e. traits that co-occur in IIS mutants or flies with increased FOXO activity. To test this hypothesis we performed qRT-PCR analysis of a major transcriptional target of FOXO, *InR*: when IIS is low, FOXO becomes active and upregulates *InR* transcription, while under high IIS FOXO is inactive and represses *InR* (Puig et al. 2003; Puig and Tjian 2005). In support of this hypothesis we found that the LL allele had a ~12% higher level of *InR* transcript than the HL allele (Fig. S8; Table S6, Supporting Information). Dietary conditions also affected *InR* levels, with flies raised on molasses producing more *InR* than flies raised on sucrose diet (Fig. S8; Table S6, Supporting Information).

## Discussion

### CONNECTING ADAPTIVE CLINAL PHENOTYPES TO GENOTYPES

Here we have studied the life-history effects of a strongly clinally varying, presumably adaptive polymorphism in the IIS gene *foxo*, a naturally segregating variant identified from our previous genomic analysis of the North American latitudinal cline (Fabian et al. 2012).

As hypothesized by de Jong and Bochdanovits (2003), genes of the IIS/TOR pathway might represent particularly promising candidates underlying clinal life-history adaptation in *D. melanogaster*: (1) laboratory mutants in this pathway often mirror life-history traits and trade-offs observed in natural populations (de Jong and Bochdanovits 2003; Clancy et al. 2001; Tatar et al. 2001; Tatar and Yin 2001; Tatar et al. 2003; Paaby et al. 2010; Flatt et al. 2013; Paaby et al. 2014; Flatt and Partridge 2018); (2) reproductive dormancy in response to cool temperature and short photoperiod, a genetically variable and clinal trait (Williams and Sokolowski 1993; Schmidt et al. 2005a; Schmidt and Conde 2006; Schmidt et al. 2005b; Schmidt and Paaby 2008), is physiologically regulated by IIS (Williams et al. 2006; Flatt et al. 2013; Kubrak et al. 2014; Schiesari et al. 2016; Zhao et al. 2016; Andreatta et al. 2018); (3) genomic analyses of clinal differentiation has identified many clinal SNPs in the IIS/TOR pathway presumably shaped by spatially varying selection (Fig. 1; Kolaczkowksi et al. 2011; Fabian et al. 2012; Kapun et al. 2016b); and (4) genome-wide analyses of variation in size-related traits have identified novel regulators of growth, several of which interact with the IIS/TOR pathway (Vonesch et al. 2016; Strassburger et al. 2017). For example, in support of the idea that variation in IIS contributes to clinal adaptation in *D. melanogaster*, Paaby and colleagues have identified a clinal indel polymorphism in *InR* with pleiotropic effects on development, body size, fecundity, lifespan, oxidative stress resistance, chill coma recovery, and insulin signaling (Paaby et al. 2010, 2014). Our results on *foxo* lend further support to the hypothesis of de Jong and Bochdanovits (2003).

### THE EFFECTS OF NATURAL VERSUS NULL ALLELES AT THE *FOXO* LOCUS

Previous work with loss-of-function mutants and transgenes has uncovered a major role of *foxo* in the regulation of growth, lifespan and resistance to starvation and oxidative stress (Jünger et al. 2003; Puig et al. 2003; Kramer et al. 2003; Giannakou et al. 2004; Hwangbo et al. 2004; Kramer et al. 2008; Slack et al. 2011), but nothing is known yet about the effects of natural alleles at this locus. An important distinction in this context is that null mutants, by definition, reveal the complete set of functions and phenotypes of a given gene and may therefore be highly pleiotropic, whereas ‘evolutionarily relevant’ mutations or alleles might have much more subtle effects, with little or no pleiotropy (Stern 2000). Based on our knowledge of the traits affected by *foxo* in null mutants and transgenes (Jünger et al. 2003; Kramer et al. 2003, 2008; Slack et al. 2011), we measured how the clinal 2-SNP variant affects size traits and starvation resistance.

Although we could not predict with certainty the directionality and/or the degree of pleiotropy of the allelic effects *a priori*, we found that the *foxo* polymorphism differentially affects several size-related traits and starvation resistance, phenotypes known to be affected by the *foxo* locus. With regard to growth and size, our findings from natural variants agree well with functional genetic studies showing that genetic manipulations of the *foxo* locus affect body size and wing area (Jünger et al. 2003; Slack et al. 2011; Tang et al. 2011). Similarly, our observation that variation at *foxo* affects survival and fat content upon starvation is consistent with the fact that *foxo* mutants display reduced starvation resistance (Jünger et al. 2003; Kramer et al. 2003, 2008). In contrast, although *foxo* null mutants produce viable adults (Jünger et al. 2003; Slack et al. 2011), whether distinct *foxo* alleles vary in viability has not yet been examined; here we find that the two natural alleles differ in egg-to-adult survival. We also asked whether the alleles differentially affect mRNA abundance of *InR*, a transcriptional target of FOXO (Puig et al. 2003; Puig and Tjian 2005). Indeed, the LL allele had higher *InR* mRNA levels, consistent with the LL genotype exhibiting reduced IIS and higher FOXO activity.

For most traits measured, both alleles reacted plastically to changes in diet and temperature in the direction predicted from previous work (Partridge et al. 1994a, 1994b; Lee and Jang 2014; Lihoreau et al. 2016; Mathur and Schmidt 2017), yet we found very little evidence for allele by environment (G × E) interactions.

While our experimental design does not allow us to disentangle the contribution of the 2 individual SNPs to the total effects seen for the *foxo* polymorphism, our results suggest that the naturally occurring alternative alleles at *foxo* we have examined here – and which are defined by only two linked SNP positions –can apparently have quite strong pleiotropic (or, via LD, correlational) effects upon multiple complex life-history traits, including on viability, several proxies of size and on starvation resistance (for estimates of allelic effect sizes see Table S4, Supporting Information). This is consistent with the pleiotropic effects seen in *foxo* loss-of-function mutant alleles (see references above) and might support the idea that the architecture of life-history traits, which are connected via multiple trade-offs, is inherently pleiotropic (Williams 1957; Finch and Rose 1995; Flatt et al. 2005; Flatt and Promislow 2007; Flatt and Schmidt 2009; Flatt et al. 2013; Paaby et al. 2014); it also provides a contrast to the model from evo-devo which posits that most evolutionarily relevant mutations should exhibit little or no pleiotropy (Stern 2011). In particular, the pleiotropic effects of the *foxo* variant might explain why this polymorphism might be maintained, through some form of balancing selection, in natural populations along the cline.

### INSULIN SIGNALING, CLINALITY, AND COUNTERGRADIENT VARIATION

How does the *foxo* variant contribute to phenotypic clines observed across latitude? High-latitude flies tend to be characterized, for example, by larger body size, decreased fecundity, longer lifespan and improved stress resistance as compared to low-latitude flies, and this differentiation is genetically based (Coyne and Beecham 1987; Schmidt et al. 2005a, 2005b; Schmidt and Paaby 2008; Mathur and Schmidt 2017; Durmaz et al. 2018). Do the allelic effects go in the same direction as the latitudinal gradient, representing cogradient variation, or do certain allelic effects run counter to the cline, representing countergradient variation (Levins 1968; Conover and Schultz 1995)? Cogradient variation occurs when diversifying selection favors different traits in different environments, as expected from selection along a cline, whereas countergradient variation occurs when stabilizing selection favors similar traits in different environments (Conover and Schultz 1995; Marcil et al. 2006).

Consistent with clinal expectation, the HL allele confers larger size (Coyne and Beecham 1987; de Jong and Bochdanovits 2003); increased wing:thorax ratio, which corresponds to reduced ‘wing loading’, a trait hypothesized to be adaptive for flight at cold temperature (Stalker 1980; David et al. 1994; Azevedo et al. 1998; Frazier et al. 2008; Bhan et al. 2014); and reduced viability (Folguera et al. 2008). Conversely, the LL allele exhibits smaller size, increased wing loading, and higher viability. Thus, the *foxo* variant contributes to the observed phenotypic cline in the predicted direction (gradient or cogradient variation) and appears to be maintained by spatially varying selection (for a remarkable example where size is subject to countergradient – not cogradient – variation along an altitudinal gradient in Puerto Rican *D. melanogaster* see Levins, 1968, 1969). Importantly, our results for size-related traits are consistent with independent assays under constant environmental conditions (Betancourt et al. 2018) and suggest a major contribution of the *foxo* polymorphism to the clinality of body size.

For starvation resistance, we found – contrary to clinal predictions – that the HL allele is less resistant than the LL allele, consistent with countergradient variation. Interestingly, a similar countergradient effect (on body size) was found for the *InR* polymorphism mentioned above: the high-latitude *InR^short^* allele confers smaller size, even though flies from high-latitude populations are normally larger (Paaby et al. 2014). Likewise, for a clinal variant of *neurofibromin 1* (*Nf1*) the high-latitude haplotype has smaller wing size, an effect that runs counter to the cline (Lee et al. 2013). However, as mentioned in the methods, we can of course not completely rule out potentially confounding LD effects that might account for this unexpected result with regard to starvation resistance.

In terms of the physiological effects of IIS, temperate fly populations might be characterized by ‘thrifty’ genotypes with high IIS, whereas tropical populations might have a higher frequency of ‘spendthrift’ genotypes with low IIS (de Jong and Bochdanovits 2003). Our finding that the low-latitude *foxo* allele likely exhibits increased FOXO activity and lower IIS seems to support this, yet Paaby et al. (2014) found that IIS was lower for the high-latitude *InR* allele. The directionality of IIS effects along the cline thus remains difficult to predict.

As noted by Lee et al. (2013) and Paaby et al. (2014), clinal variants subject to countergradient effects might interact epistatically with other loci affecting the trait, or they might be affected by antagonistic selection pressures (Schluter et al. 1991). Conflicting selection pressures on clinal variants might be particularly acute when they exhibit pleiotropic effects on multiple traits, as is the case for the polymorphisms at *Nf1, InR*, and *foxo*. These examples illustrate the complexity of dissecting clinal selection and the genotype-phenotype map underlying clinal adaptation (Lee et al. 2013; Paaby et al. 2014; Flatt 2016).

With regard to starvation resistance, a caveat is that we found the LL allele to be more resistant, while Betancourt et al. (2018) found the HL allele to be more resistant. This discrepancy might be due to differences in assay protocols: Betancourt et al. (2018) did not use agar which could impose some desiccation in addition to starvation stress. Interestingly, desiccation resistance is known to vary latitudinally along the North America east coast (Rajpurohit et al. 2018), but whether the *foxo* polymorphism examined here affects survival upon desiccation remains unknown and awaits future study.

### GROWING EVIDENCE FOR A ROLE OF IIS IN LIFE-HISTORY ADAPTATION

The IIS pathway provides an excellent example of how mechanistic and evolutionary insights might be combined to gain a more complete understanding of the ultimate and proximate determinants of life-history adaptation (Finch and Rose 1995; Houle 2001; Flatt and Heyland 2011). Since the 1990s, a great deal has been learned about the genetic, developmental and physiological effects of this pathway in model organisms. This work has shown that IIS mutants affect major fitness-related traits, and this in turn has illuminated our understanding of the molecular underpinnings of growth, size, lifespan and trade-offs (Partridge and Gems 2002; Tatar et al. 2003; Flatt et al. 2005; Flatt and Heyland 2011; Flatt et al. 2013). In particular, these studies have revealed that the IIS pathway plays an evolutionarily conserved role in the physiological regulation of longevity (Partridge and Gems 2002; Tatar et al. 2003); they have also given us some of the clearest examples of alleles exhibiting antagonistic pleiotropy (Williams 1957; Flatt and Promislow 2007; and references above).

The functional characterization of this pathway therefore promised an opportunity for evolutionary geneticists to identify natural variants involved in life-history evolution (de Jong and Bochdanovits 2003). Yet, ‘life history loci’ identified via functional genetic analysis need not necessarily contribute to standing variation for these traits in the wild (Flatt 2004; Flatt and Schmidt 2009; Fabian et al. 2018). For some time, it thus remained unclear whether natural variation in this pathway impacts variation in fitness-related traits in natural populations (see Reznick 2005; Fabian et al. 2018).

Today, we have growing evidence that variation in IIS indeed can make an important contribution to life-history variation in flies and other insects, worms, fish, reptiles and mammals, including effects on longevity in humans (e.g., de Jong and Bochdanovits 2003; Williams et al. 2006; Flachsbart et al. 2008; Suh et al. 2008; Willcox et al. 2008; Alvarez-Ponce et al. 2009; Sparkman et al. 2009, 2010; Paaby et al. 2010; Stuart and Page 2010; Dantzer and Swanson 2012; Jovelin et al. 2014; Paaby et al. 2014; Swanson and Dantzer 2014; McGaugh et al. 2015; Schwartz and Bronikowski 2016; Zhao et al. 2016; and references therein). On the other hand, ‘evolve and resequence’ studies of *Drosophila* longevity have failed to find a major contribution of standing variation in IIS to evolved changes in life history and lifespan, perhaps suggesting that the IIS pathway might be selectively constrained, at least with regard to the evolution of certain traits (e.g., Remolina et al. 2012; Fabian et al. 2018; Flatt and Partridge 2018). In sum, this body of work illustrates how one might be able to connect genotypes to molecular mechanisms to components of fitness by studying a fundamentally important physiological pathway from multiple angles (Finch and Rose 1995; Houle 2001; Flatt and Heyland 2011; Flatt et al. 2013).

## Conclusions

Here we have found that a strongly clinal polymorphism at the *foxo* locus (which might be viewed as a marker for functionally significant alleles) has effects on several fitness components known to vary across latitude, including viability, size-related traits, starvation resistance and fat content. The directionality of most of these effects matches overall phenotypic clines observed along the North American east coast, especially with regard to size (e.g., Coyne and Beecham 1987; Schmidt et al. 2005a, 2005b; Schmidt and Paaby 2008; Durmaz et al. 2018). Together, our results thus suggest that standing variation in the IIS pathway makes an important and – at least partly – predictable contribution to life-history clines in *Drosophila*.

## ACKNOWLEDGEMENTS

We thank Jeff Jensen and two anonymous reviewers for helpful comments on our paper; the members of the Flatt and Schmidt labs for assistance in the lab; and Fisun Hamaratoglu, Tad Kawecki, Wolf Blanckenhorn and Marc Tatar for insightful discussion and/or comments on an early version of our manuscript. Our research was supported by the Swiss National Science Foundation (SNSF grants PP00P3_133641; PP00P3_165836; 310030E-164207; 310003A182262 to TF), the Austrian Science Foundation (FWF P21498-B11 to TF), the National Institutes of Health (NIH R01GM100366 to PS), the National Science Foundation (NSF DEB 0921307 to PSS), and the Department of Ecology and Evolution at the University of Lausanne. Parts of this paper were written while TF was a Visiting Professor in the Research Training Group 2200 ‘Evolutionary Processes in Adaptation and Disease’ at the Institute for Evolution and Biodiversity, University of Münster, Germany, and supported by Mercator Fellowship from the German Research Foundation (DFG) to TF.

## DATA ACCESSIBILITY

Phenotypic raw data are available from Dryad at doi to be added upon acceptance.

## AUTHOR CONTRIBUTIONS

T.F. and P.S. conceived the project. D.F. and M.K. identified the *foxo* SNPs and performed genomic analyses. T.F., P.S., E.D. and S.R. designed the experiments. SR and NB established reconstituted outbred populations. E.D., S.R. and N.B. performed the experiments. E.D., N.B., P.S. and T.F. analyzed the data. E.D., P.S. and T.F. wrote the paper with input from the other authors.

## Supporting Information

**Figure S1.**
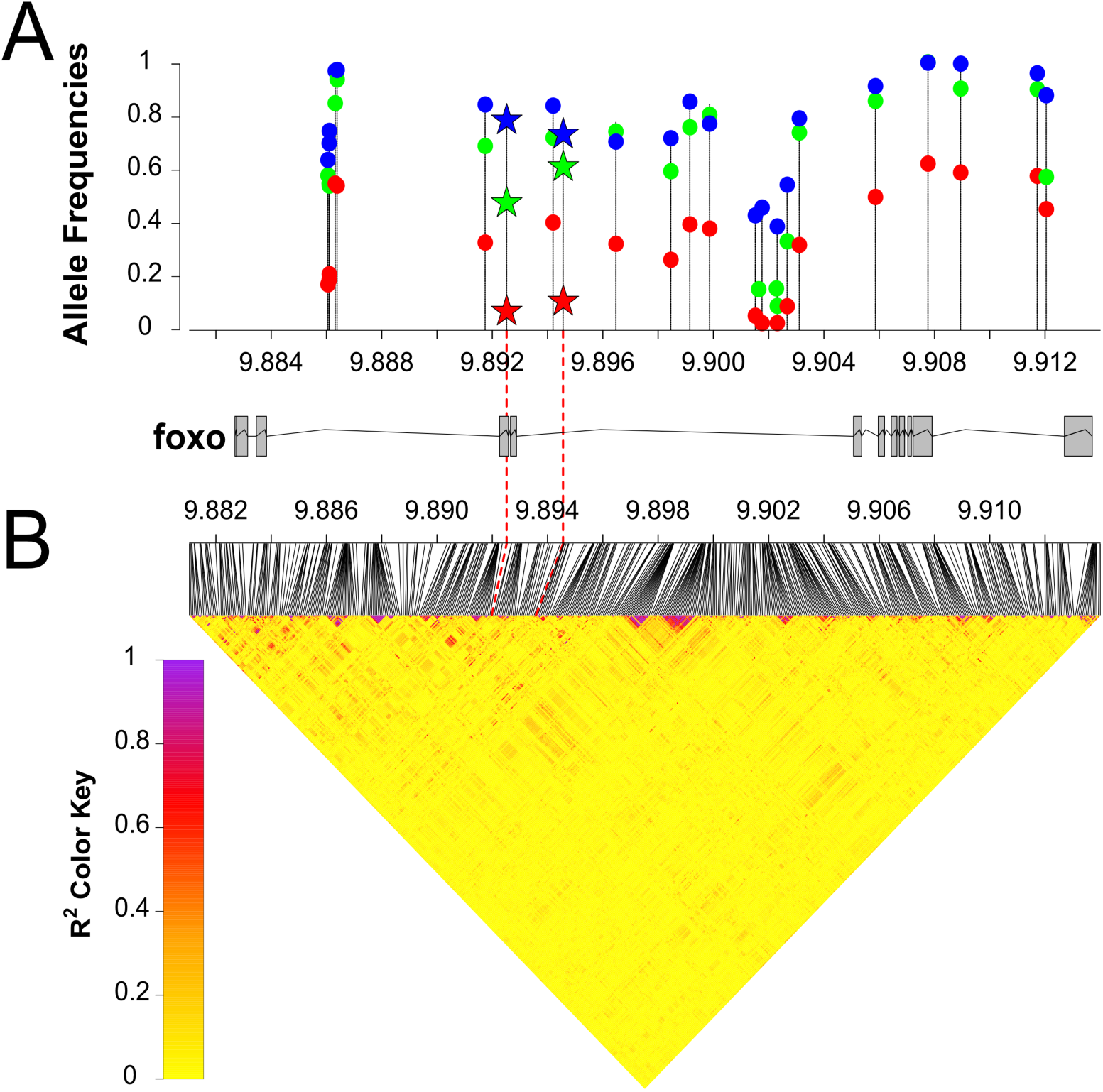
Clinal *foxo* candidate SNPs. (A) Allele frequencies of clinal *foxo* SNPs in Florida (red), Pennsylvania (green) and Maine (blue), identified by Fabian et al. (2012) and conditioned to raise in frequency from Florida to Maine. The two strongly clinal *foxo* SNPs studied here are marked with star symbols. Note that the SNP in-between the two focal SNPs is much less strongly clinal, with a much higher frequency in Florida than the 2 candidate SNPs. The x-axis shows the genomic position of the SNPs on chromosome *3R* in million base pairs (Mbp). The plot underneath the x-axis shows the gene model for *foxo*. (B) Linkage disequilibrium (LD; as measured by pairwise *r*^2^) among all polymorphic *foxo* SNPs (minor allele frequency ≥ 0.1) in the DGRP lines used to set up experimental populations (see Materials and Methods section). The two focal SNPs are in perfect LD in the experimental populations (*r*^2^ =1), but there is no significant LD among other, non-focal sites. Nonetheless, we cannot rule out with certainty that other SNPs are in LD with our two focal SNPs; a cautious interpretation would thus be to view our focal SNPs as representing “tag SNPs”. Also see Fig. S3; also see analyses in Betancourt et al. (2018).

**Figure S2.**
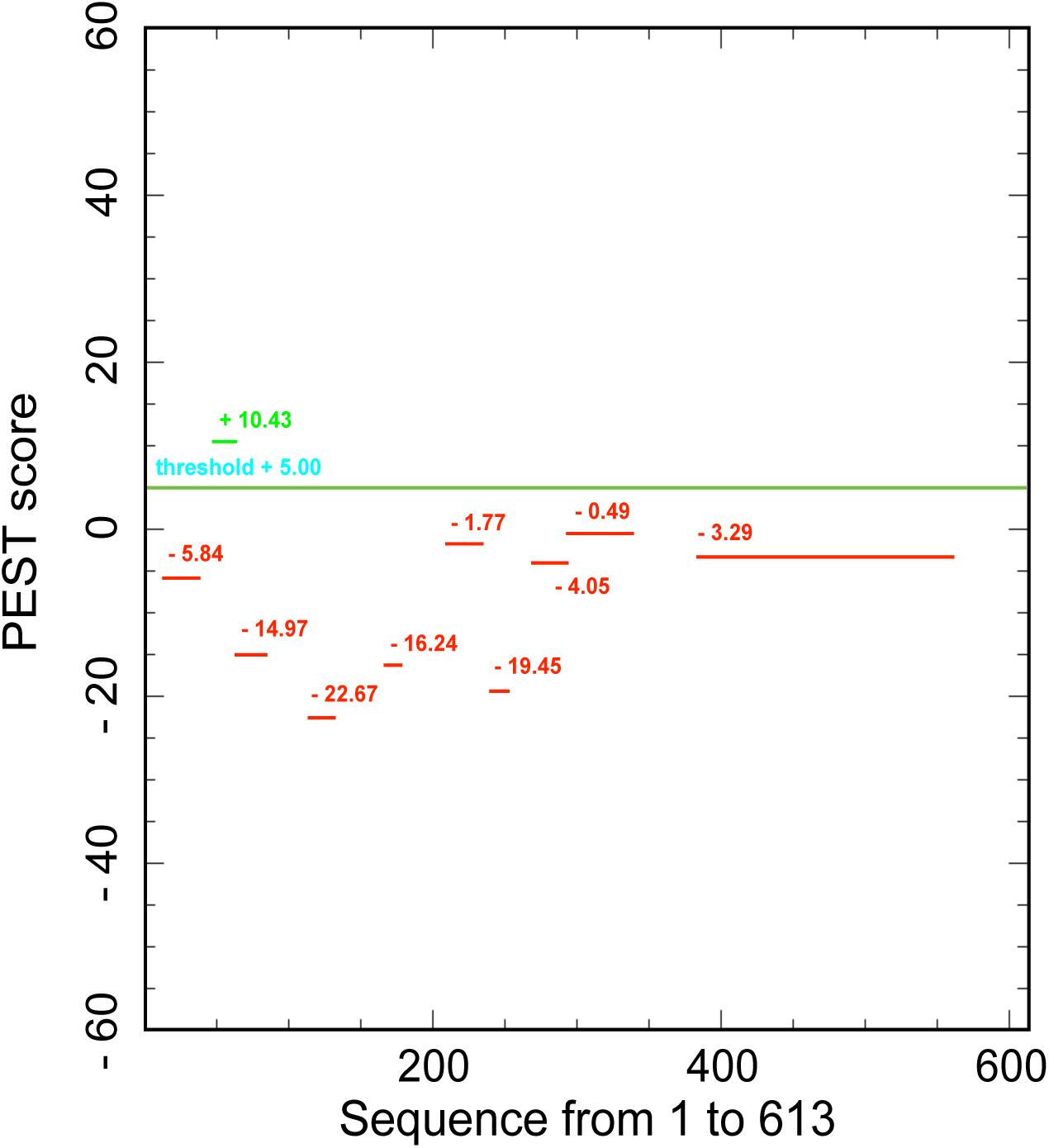
PEST motif prediction for FOXO. The T/G polymorphism in *foxo* at position *3R*: 9894559, is predicted to be located in the PEST region of the FOXO protein (analysis of *foxo* sequence using ExPASy [Artimo et al., 2012]); PEST motifs serve as protein degradation signals (Artimo et al., 2012). The potential PEST motif (RPENFVEPTDELDSTK) between amino acid positions 49 and 64 (shown in green) encompasses the *foxo* SNP at position 51 (E = glutamic acid).

**Figure S3.**
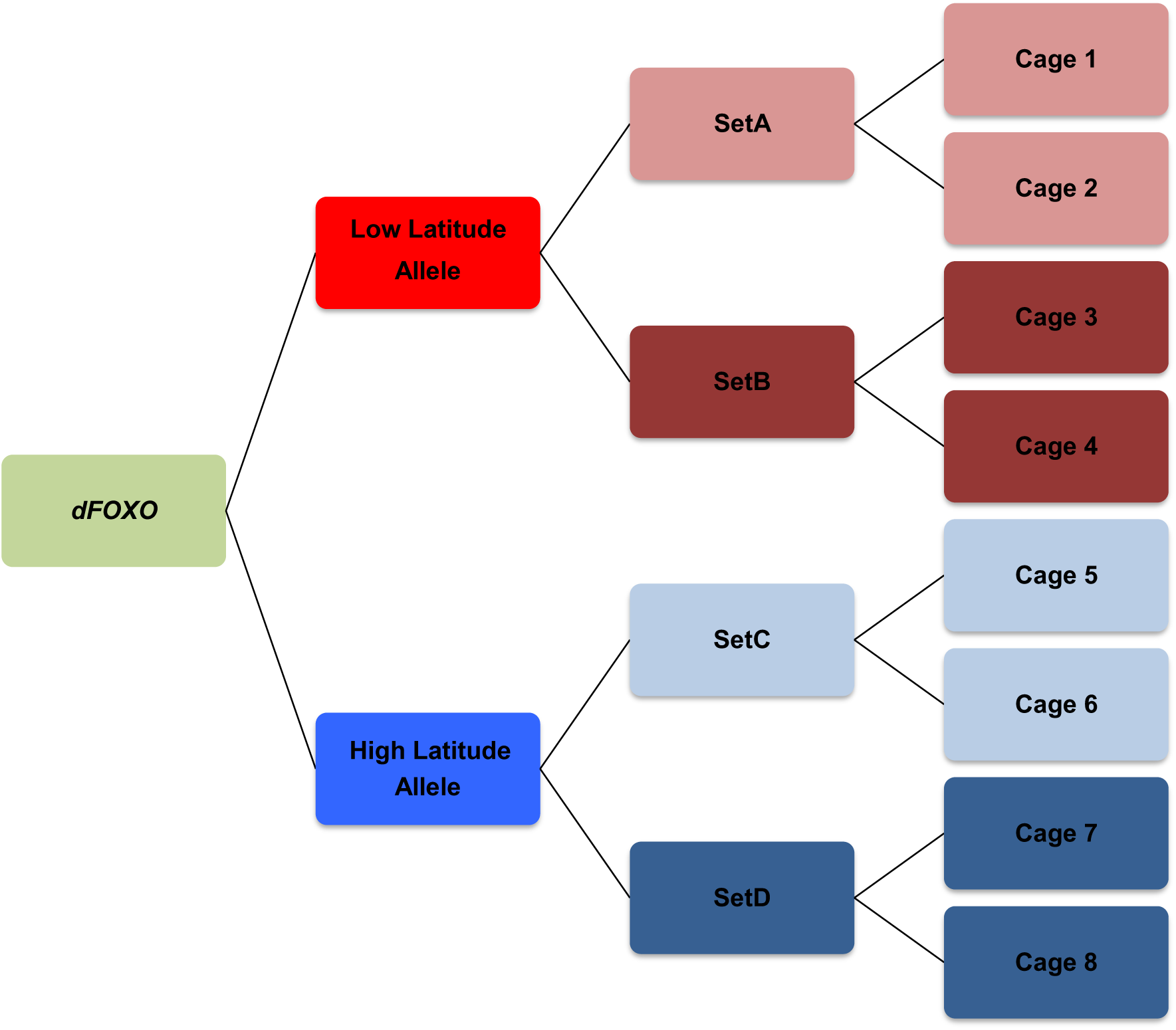
Experimental design for reconstituted outbred *foxo* populations. We isolated the 2-SNP *foxo* variant by reconstituting outbred populations, fixed for either the low- or high-latitude allele, from lines of the *Drosophila* Genetic Reference Panel (DGRP). Each *foxo* allele was represented by two independent sets of distinct DGRP lines, with two replicate cages per set. See Materials and Methods section for details; also see Fig. S1B; also see analyses in Betancourt et al. (2018).

**Figure S4.**
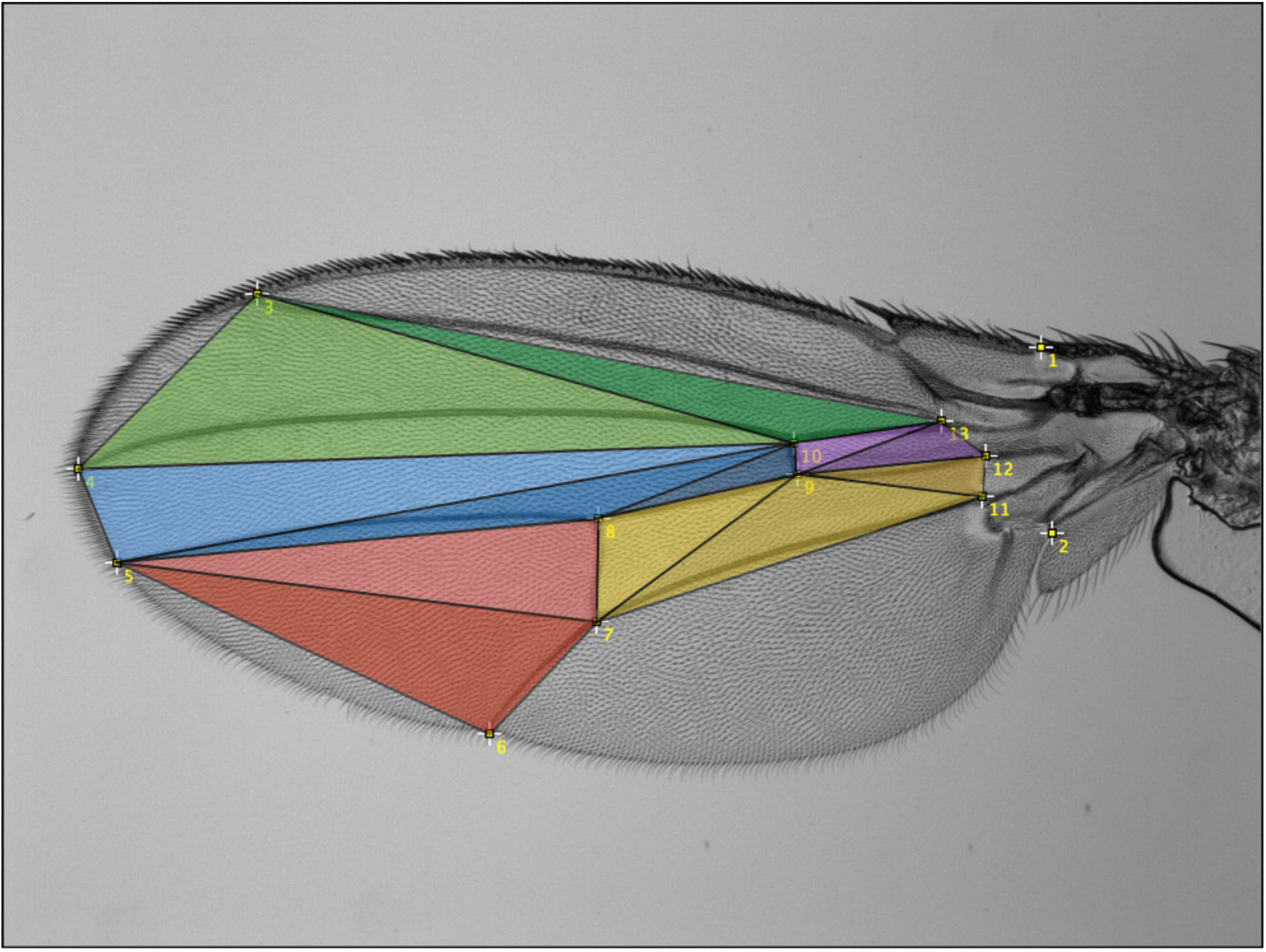
Coordinates of landmarks used to estimate wing area. We calculated the total wing area encompassed by 12 landmarks (in yellow) by splitting the polygon up into triangles (shown in different colors) and by summing across the areas defined by these triangles. See Materials and Methods section for details.

**Figure S5.**
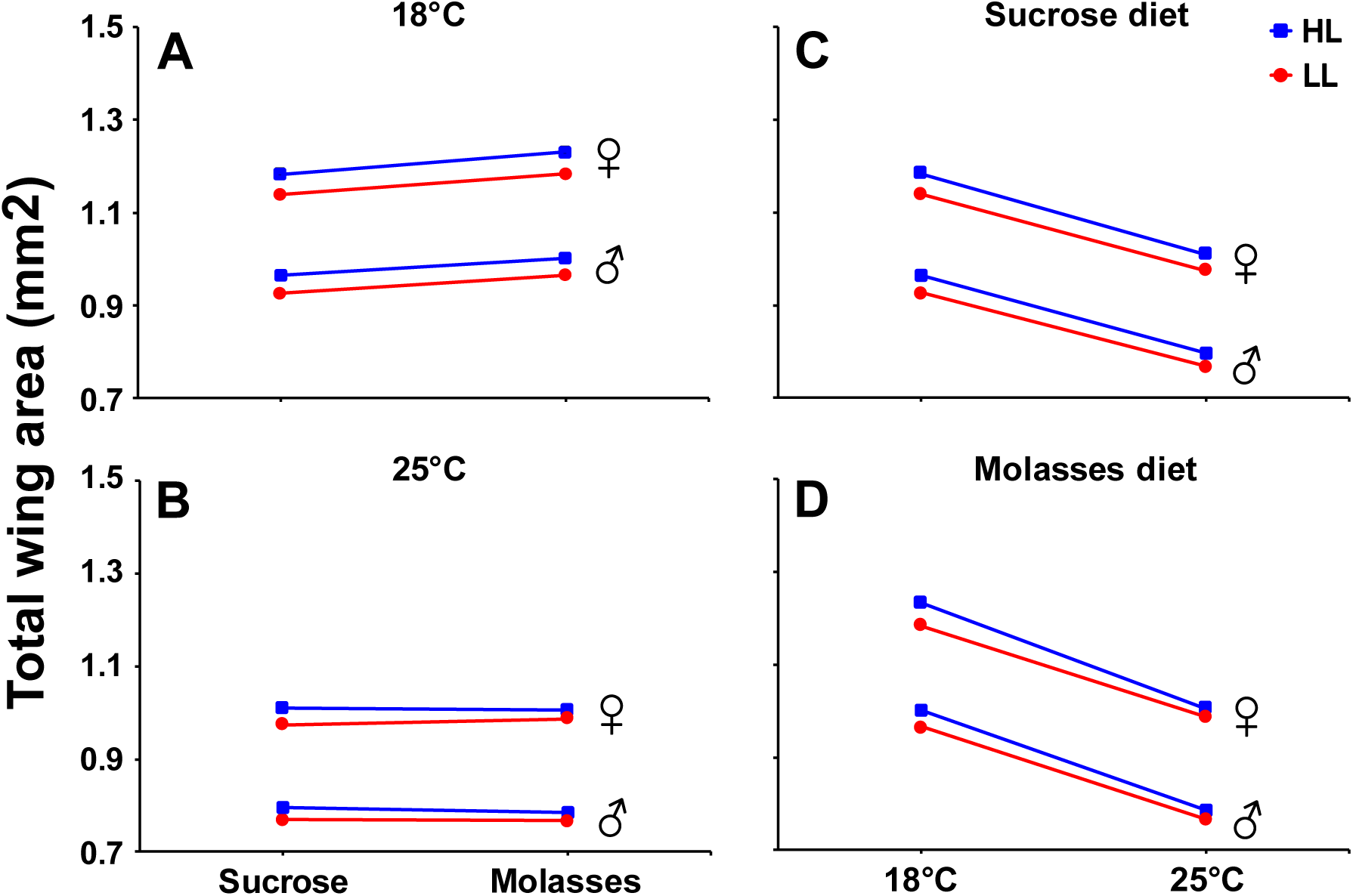
Effects of the *foxo* variant on total wing area. Effects of the clinal *foxo* variant on wing area (mm^2^) in females and males. (A) Dietary reaction norms at 18°C. (B) Dietary reaction norms at 25°C. (C) Thermal reaction norms on sucrose diet. (D) Thermal reaction norms on molasses diet. Shown are means and standard errors. Red lines: low-latitude (LL) allele, blue lines: high-latitude (HL) allele. See Results section for details.

**Figure S6.**
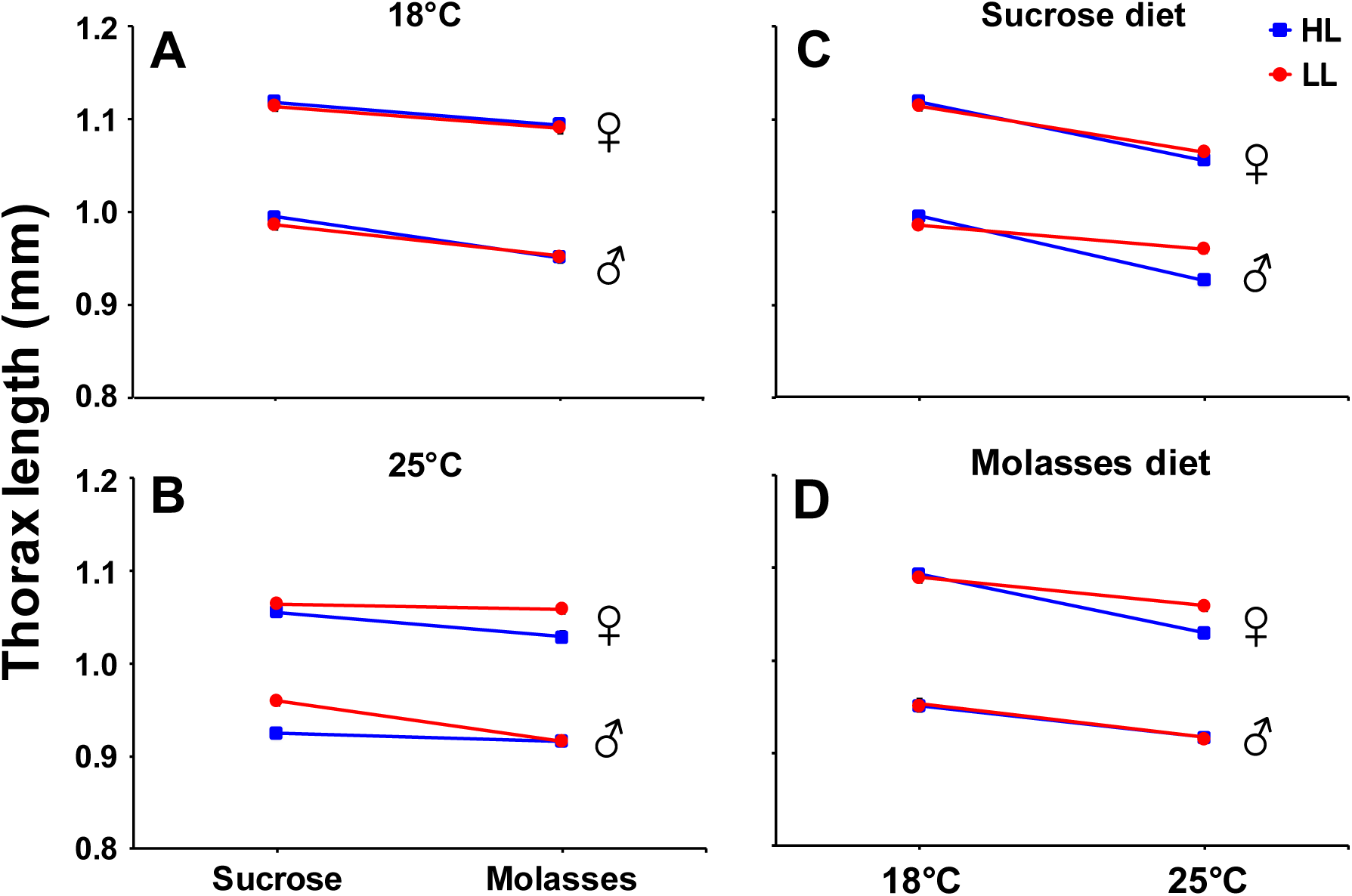
Effects of the *foxo* variant on thorax length. Effects of the clinal *foxo* variant on thorax length (mm) in females and males. (A) Dietary reaction norms at 18°C. (B) Dietary reaction norms at 25°C. (C) Thermal reaction norms on sucrose diet. (D) Thermal reaction norms on molasses diet. Shown are means and standard errors. Red lines: low-latitude (LL) allele, blue lines: high-latitude (HL) allele. See Results section for details.

**Figure S7.**
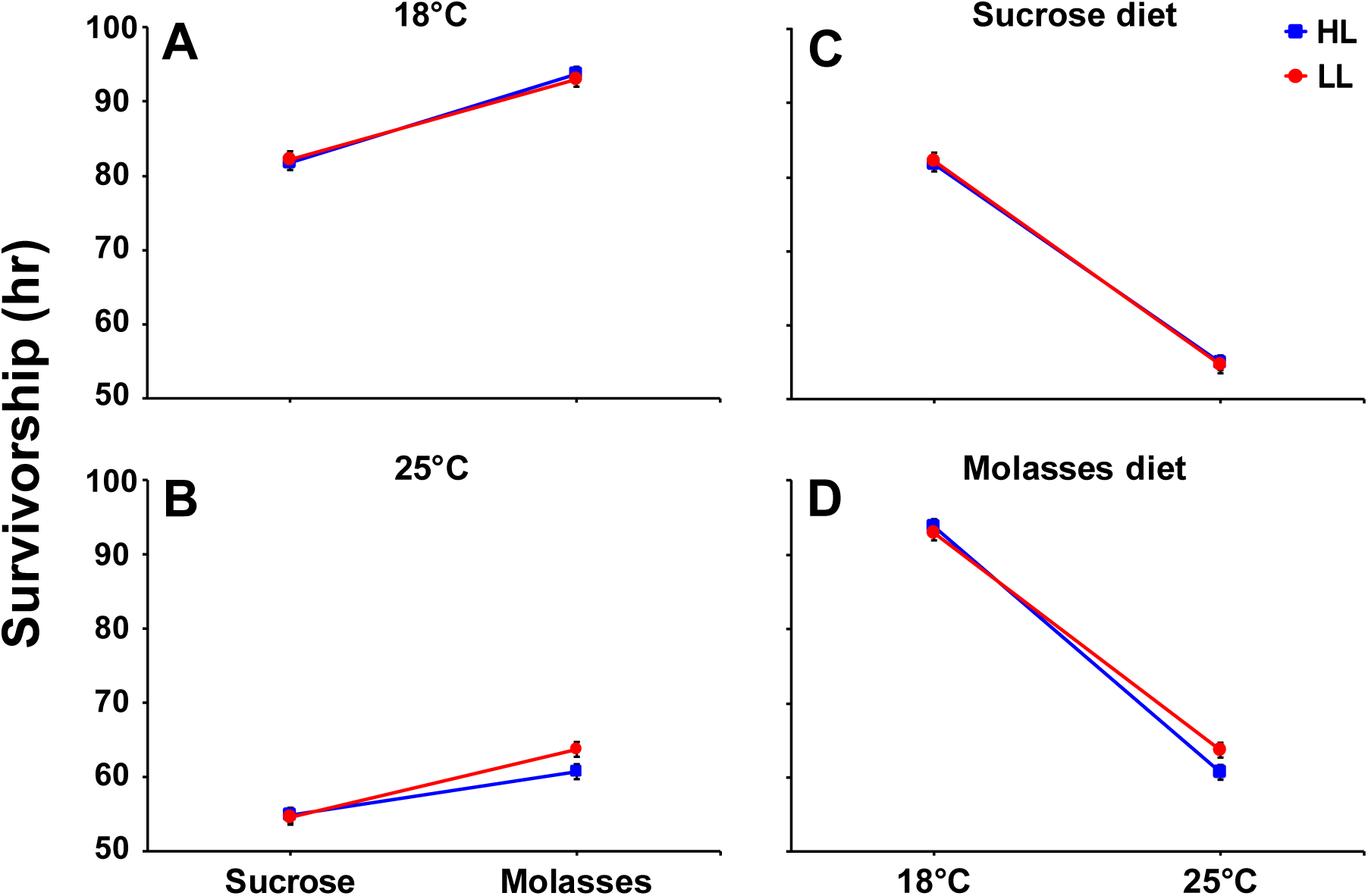
Effects of the *foxo* variant on male survival upon starvation. Effects of the clinal *foxo* polymorphism on the duration of survival (in hrs) upon starvation in males. (A) Dietary reaction norms at 18°C. (B) Dietary reaction norms at 25°C. (C) Thermal reaction norms on sucrose diet. (D) Thermal reaction norms on molasses diet. Shown are means and standard errors. Red lines: low-latitude (LL) allele, blue lines: high-latitude (HL) allele. See Results section for details.

**Figure S8.**
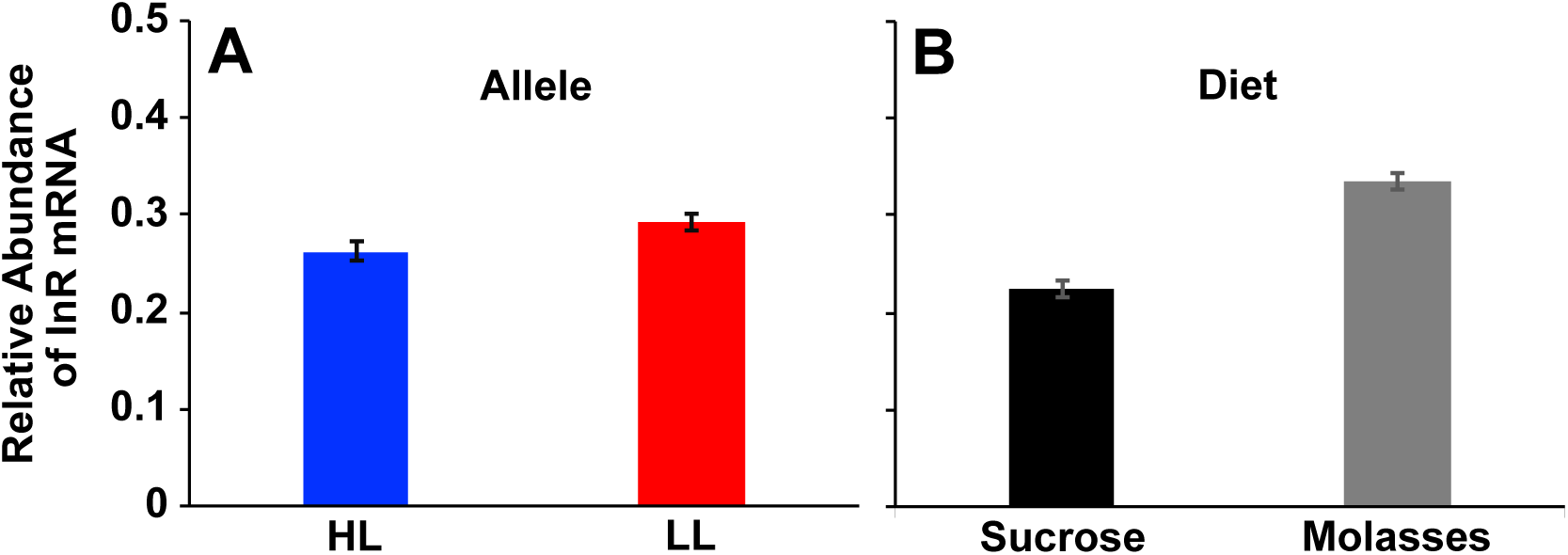
Effects of the *foxo* variant on relative abundance of insulin-like receptor (InR) transcription levels. (A) Low-latitude (LL) allele has higher level of InR transcription than the high-latitude (HL) allele. (B) Carbohydrate-rich molasses diet resulted in more InR transcripts than the sucrose diet. Shown are means and standard errors. See Results section for details.

**Table S1.** Details of design of reconstituted outbred population cages. HL: high-latitude *foxo* allele; LL: low-latitude *foxo* allele. See Materials and Methods section for details.

| Allele | Position | Set | Cage number | DGRP lines |
| --- | --- | --- | --- | --- |
| LL | 3R:9892517 + 9894559<br>(GG) | A | 1 | 26, 57, 73, 75, 91, 101, 105, 161, 176, 280,<br>313, 318, 367, 371, 375, 377, 378, 379 |
| LL | 3R:9892517 + 9894559<br>(GG) | A | 2 | 26, 57, 73, 75, 91, 101, 105, 161, 176, 280,<br>313, 318, 367, 371, 375, 377, 378, 379 |
| LL | 3R:9892517 + 9894559<br>(GG) | B | 3 | 208, 373, 406, 426, 440, 491, 492, 508, 513,<br>535, 639, 646, 757, 761, 796, 805, 812, 852 |
| LL | 3R:9892517 + 9894559<br>(GG) | B | 4 | 208, 373, 406, 426, 440, 491, 492, 508, 513,<br>535, 639, 646, 757, 761, 796, 805, 812, 852 |
| HL | 3R:9892517 + 9894559<br>(AT) | C | 5 | 40, 41, 42, 69, 83, 109, 142, 153, 158, 177,<br>195, 229, 233, 365, 370, 380, 391, 405 |
| HL | 3R:9892517 + 9894559<br>(AT) | C | 6 | 40, 41, 42, 69, 83, 109, 142, 153, 158, 177,<br>195, 229, 233, 365, 370, 380, 391, 405 |
| HL | 3R:9892517 + 9894559<br>(AT) | D | 7 | 45, 332, 338, 443, 517, 531, 595, 703, 705,<br>707, 774, 790, 804, 820, 837, 855, 879, 890 |
| HL | 3R:9892517 + 9894559<br>(AT) | D | 8 | 45, 332, 338, 443, 517, 531, 595, 703, 705,<br>707, 774, 790, 804, 820, 837, 855, 879, 890 |

**Table S2.** Nutritional value and composition of sucrose and molasses diets. Table S2a: nutritional values of fly food ingredients per 100 g; Table S2b: recipe for sucrose and molasses diets; Table S2c: comparison of nutritional values of sucrose and molasses diets. See Materials and Methods section for details. The sucrose diet is the standard medium used in our laboratory in Lausanne; the recipe for the molasses diet follows that recipe of the Bloomington *Drosophila* Stock Center (BDSC) but uses different products for the food ingredients. The principal (but not exclusive) differences between the two diets are their carbohydrate source (sucrose vs. molasses) and their protein:carbohydrate (P:C) ratios.

| **S2a. Nutritional values of ingredients in 100g of fly food** |  |  |  |  |
| --- | --- | --- | --- | --- |
|  | **Yeast** | **Cornmeal** | **Sucrose** | **Molasses** |
| Energy (kcal) | 310 | 345 | 400 | 290 |
| Protein (g) | 45 | 8 | 0 | 0 |
| Total carbohydrates (g) | 15 | 74 | 100 | 75 |

| **S2b. Food recipes for sucrose and molasses diets** |  |  |
| --- | --- | --- |
|  | **Sucrose** | **Molasses** |
| Cornmeal (g/L) (*Polenta, Migros*) | 50 | 61.3 |
| Yeast (g/L) (*Actilife, Migros*) | 50 | 12.4 |
| Sugar (g/L) (*Cristal, Migros*) | 50 | 0 |
| Molasses (g/L) (*Zuckerrohrmelasse, EM Schweiz*) | 0 | 109.6 |
| Agar (g/L) (*Drosophila Agar Type II, Genesee*) | 7 | 6 |
| Nipagin 10% (ml/L) (*Sigma Aldrich*) | 10 | 14.3 |
| Propionic acid (ml/L) (*Sigma Aldrich*) | 6 | 6 |

| **S2c. Nutritional values of sucrose and molasses diets** |  |  |
| --- | --- | --- |
|  | **Sucrose** | **Molasses** |
| Energy (kcal) | 527.50 | 567.77 |
| Protein (g/L) | 26.50 | 10.48 |
| Total carbohydrate (g/L) | 94.50 | 129.42 |
| P:C ratio | ~ 1:3.6<br>($\approx 0.28$) | ~ 1:12.3<br>($\approx 0.08$) |

**Table S3.** Summary of effect size estimates (Cohen’s *d*) for viability, femur length, wing area, thorax length, starvation resistance, and fat (TAG) content. White and grey cells show results for females and males, respectively. *d* = 0.01, very small; *d* = 0.20, small; *d* = 0.50, medium; *d* = 0.80, large; *d* = 1.20, very large.

| Factor | 18°C<br>Sucrose diet | 18°C<br>Molasses diet | 25°C<br>Sucrose diet | 25°C<br>Molasses diet |
| --- | --- | --- | --- | --- |
| Viability | 0.49 | 0.50 | 0.54 | 0.89 |
| Femur Length | 0.09 | 0.17 | 0.49 | 0.00 |
|  | 0.25 | 0.05 | 0.14 | 0.20 |
| Wing Area | 0.59 | 0.67 | 0.66 | 0.35 |
|  | 0.68 | 0.62 | 0.72 | 0.48 |
| Thorax Length | 0.13 | 0.08 | 0.20 | 0.81 |
|  | 0.26 | 0.07 | 1.15 | 0.00 |
| Starvation<br>Resistance | 0.11 | 0.34 | 0.24 | 0.51 |
|  | 0.03 | 0.04 | 0.03 | 0.26 |
| TAG content<br>(Fed) | 0.19 | 0.70 | 0.25 | 0.72 |
| TAG content<br>(Starved) | 0.72 | 0.24 | 0.04 | 0.04 |

**Table S4.** Summary of ANOVA results for wing area, thorax length, and male starvation resistance (also cf. Table S5). White and grey cells show the results for females and males, respectively; data for starvation resistance are for males only. * *p* < 0.05; ** *p* < 0.01; *** *p* < 0.001. See Results section for details.

| Factor in ANOVA | Total Wing Area | Thorax Length | Starvation Resistance |
| --- | --- | --- | --- |
| Allele | $F_{1,32}=105.39^{***}$ | $F_{1,32}=4.33^*$ | $F_{1,32}=0.70$ |
| | $F_{1,32}=103.87^{***}$ | $F_{1,32}=3.78$ | |
| Temperature | $F_{1,912}=2852.52^{***}$ | $F_{1,422}=216.46^{***}$ | $F_{1,1553}=1711.77^{***}$ |
| | $F_{1,918}=3962.67^{***}$ | $F_{1,381}=145.46^{***}$ | |
| Diet | $F_{1,912}=48.36^{***}$ | $F_{1,422}=31.90^{***}$ | $F_{1,1553}=176.44^{***}$ |
| | $F_{1,918}=28.15^{***}$ | $F_{1,381}=88.62^{***}$ | |
| Allele x Temperature | $F_{1,912}=7.15^{**}$ | $F_{1,422}=10.66^{**}$ | $F_{1,1553}=0.58$ |
| | $F_{1,918}=5.89^*$ | $F_{1,381}=8.72^{**}$ | |
| Temperature x Diet | $F_{1,912}=35.96^{***}$ | $F_{1,422}=1.67$ | $F_{1,1553}=7.51^{**}$ |
| | $F_{1,918}=56.66^{***}$ | $F_{1,381}=3.48$ | |
| Allele x Diet | $F_{1,912}=0.73$ | $F_{1,422}=2.44$ | $F_{1,1553}=0.58^{***}$ |
| | $F_{1,918}=1.08$ | $F_{1,381}=2.46$ | |
| Allele x Temperature x Diet | $F_{1,912}=1.79$ | $F_{1,422}=1.89$ | $F_{1,1553}=2.48$ |
| | $F_{1,918}=0.22$ | $F_{1,381}=11.19^{***}$ | |
| Set (Allele) | $F_{2,32}=53.59^{***}$ | $F_{2,32}=8.05^{***}$ | $F_{2,32}=1.01$ |
| | $F_{2,32}=30.53^{***}$ | $F_{2,32}=7.56^{***}$ | |
| Cage (Set, Allele) | $F_{4,32}=64.45^{***}$ | $F_{4,32}=3.41^{**}$ | $F_{4,32}=12.78^{***}$ |
| | $F_{4,32}=29.58^{***}$ | $F_{4,32}=0.73$ | |

**Table S5.** Summary of REML variance component estimates for starvation resistance. White and grey cells show results for females and males, respectively.

| Random Effect | Variance Ratio | Variance Component | Std Error | 95% Lower | 95% Upper | Wald *p*-Value | Percentage of Total |
| --- | --- | --- | --- | --- | --- | --- | --- |
| **Vial(Cage,Set,Allele)** | 0.00 | -0.19 | 2.96 | -6.00 | 5.62 | 0.95 | 0.00 |
|  | 0.00 | 0.13 | 1.29 | -2.39 | 2.65 | 0.92 | 0.07 |
| **Residual** |  | 474.90 | 17.08 | 443.13 | 510.23 |  | 100.00 |
|  |  | 199.07 | 7.14 | 185.78 | 213.85 |  | 99.93 |
| **Total** |  | 474.90 | 17.08 | 443.13 | 510.23 |  | 100.00 |
|  |  | 199.21 | 7.08 | 186.03 | 213.85 |  | 100.00 |

**Table S6.**
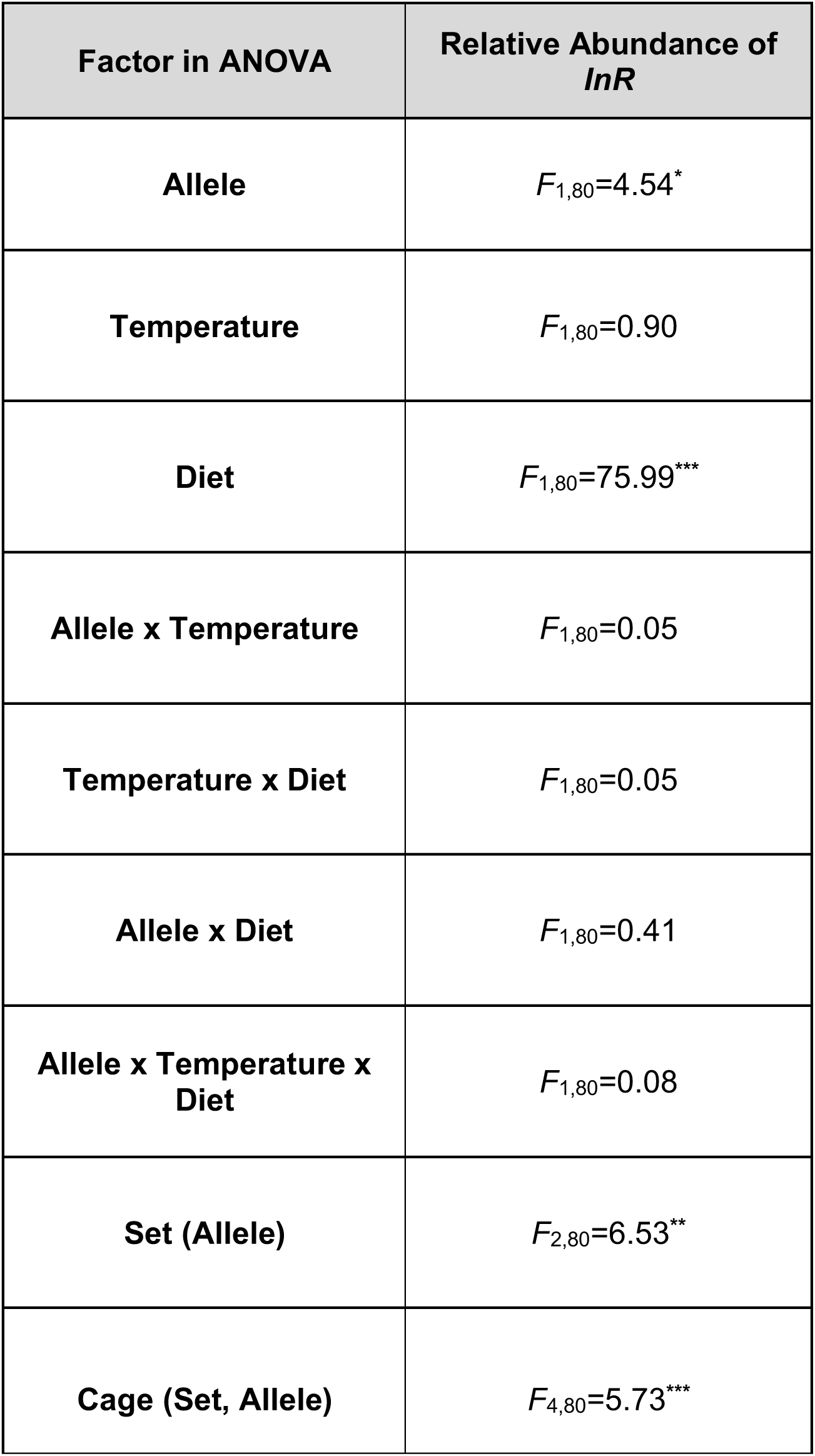
Summary of ANOVA results for relative abundance of *insulin-like receptor* (*InR*) transcript levels. * *p* < 0.05; ** *p* < 0.01; *** *p* < 0.001.

| Factor in ANOVA | Relative Abundance of *InR* |
| --- | --- |
| Allele | $F_{1,80}=4.54^*$ |
| Temperature | $F_{1,80}=0.90$ |
| Diet | $F_{1,80}=75.99^{***}$ |
| Allele x Temperature | $F_{1,80}=0.05$ |
| Temperature x Diet | $F_{1,80}=0.05$ |
| Allele x Diet | $F_{1,80}=0.41$ |
| Allele x Temperature x Diet | $F_{1,80}=0.08$ |
| Set (Allele) | $F_{2,80}=6.53^{**}$ |
| Cage (Set, Allele) | $F_{4,80}=5.73^{***}$ |

## LITERATURE CITED

Ackermann, M., Bijlsma, R., James, A. C., Partridge, L., Zwaan, B. J., and S. C. Stearns 2001. Effects of assay conditions in life-history experiments with *Drosophila melanogaster*. J. Evol. Biol. 14:199–209.

Adrion, J.R., Hahn, M. W., and B. S. Cooper. 2015. Revisiting classic clines in *Drosophila melanogaster* in the age of genomics. Trends Genet. 31:434–444.

Alvarez-Ponce, D., Aguadé, M., and J. Rozas. 2009. Network-level molecular evolutionary analysis of the insulin/TOR signal transduction pathway across 12 *Drosophila* genomes. Genome Res. 19:234–242.

Andreatta, G., Kyriacou, C. P., Flatt, T., and R. Costa. 2018. Aminergic Signaling Controls Ovarian Dormancy in *Drosophila*. Sci. Rep. 8:2030.

Artimo, P., Jonnalagedda, M., Arnold, K., Baratin, D., Csardi, G., de Castro, E., Duvaud, S., Flegel, V., Fortier, A., Gasteiger, E., Grosdidier, A., Hernandez, C., Ioannidis, V., Kuznetsov, D., Liechti, R., Moretti, S., Mostaguir, K., Redaschi, N., Rossier, G., Xenarios, I., and H. Stockinger. 2012. ExPASy: SIB bioinformatics resource portal. Nucleic Acids Res. 40:W597–W603.

Attrill, H., Falls, K., Goodman, J. L., Millburn, G. H., Antonazzo, G., Rey, A. J., Marygold, S. J., and the FlyBase Consortium. 2016. FlyBase: establishing a Gene Group resource for *Drosophila melanogaster*. Nucleic Acids Res. 44:D786–D792.

Azevedo, R. B. R., James, A. C., McCabe, J., and L. Partridge. 1998. Latitudinal variation of wing: thorax size ratio and wing-aspect ratio in *Drosophila melanogaster*. Evolution 52:1353–1362.

Barrett, R. D. H., and H. E. Hoekstra. 2011. Molecular spandrels: tests of adaptation at the genetic level. Nature Rev. Genet. 12:767–780.

Barson, N. J., Aykanat, T., Hindar, K., Baranski, M., Bolstad, G. H., Fiske, P., Jacq, C., Jensen, A. J., Johnston, S. E., Karlsson, S., Kent, M., Moen, T., Niemelä, E., Nome, T., Næsje, T. F., Orell, P., Romakkaniemi, A., Sægrov, H., Urdal, K., Erkinaro, J., Lien, S., and C. R. Primmer. 2015. Sex-dependent dominance at a single locus maintains variation in age at maturity in salmon. Nature 528:405–408.

Bergland, A. O., Behrman, E. L., O’Brien, K. R., Schmidt, P. S., and D. A. Petrov. 2014. Genomic evidence of rapid and stable adaptive oscillations over seasonal time scales in *Drosophila*. PLoS Genet. 10:e1004775.

Bergland, A. O., Tobler, R., González, J., Schmidt, P., and D. A. Petrov. 2016. Secondary contact and local adaptation contribute to genome-wide patterns of clinal variation *in Drosophila melanogaster*. Mol. Ecol. 25:1157–1174.

Behrman, E. L., Howick, V. M., Kapun, M., Staubach, F., Bergland, A. O., Petrov, D. A., Lazzaro, B. P., and P. S. Schmidt. 2018. Rapid seasonal evolution in innate immunity of wild *Drosophila melanogaster*. Proc. Roy. Soc. London B 285:20172599.

Betancourt, N. J., Rajpuorhit, S., Durmaz, E., Kapun, M., Fabian, D.K., Flatt, T., and P. S. Schmidt. 2018. Allelic polymorphism at *foxo* contributes to local adaptation in *Drosophila melanogaster*. Preprint, bioRxiv, doi: https://doi.org/10.1101/471565

Bhan, V., Parkash, R., and D. D. Aggarwal. 2014. Effects of body-size variation on flight-related traits in latitudinal populations of *Drosophila melanogaster*. J. Genet. 93:103–112.

Birney, E. 2016. The Mighty Fruit Fly Moves into Outbred Genetics. PLoS Genet. 12: e1006388.

Bochdanovits, Z., and G. de Jong. 2003a. Temperature dependence of fitness components in geographical populations of *Drosophila melanogaster*: changing the association between size and fitness. Biol. J. Linnean Soc. 80:717–725.

Bochdanovits, Z., and G. de Jong. 2003b. Experimental evolution in *Drosophila melanogaster*: interaction of temperature and food quality selection regimes. Evolution 57:1829–1836.

Böhni, R., Riesgo-Escovar, J., Oldham, S., Brogiolo, W., Stocker, H., Andruss, B. F., Beckingham, K., and E. Hafen. 1999. Autonomous control of cell and organ size by CHICO, a *Drosophila* homolog of vertebrate IRS1-4. Cell 97:865–875.

Britton, J., Lockwood, W., Li, L., Cohen, S. M., and B. A. Edgar. 2002. *Drosophila*’s insulin/PI3-kinase pathway coordinates cellular metabolism with nutritional conditions. Dev. Cell 2:239–249.

Brogiolo, W., Stocker, H., Ikeya, T., Rintelen, F., Fernandez, R., and E. Hafen. 2001. An evolutionarily conserved function of the *Drosophila* insulin receptor and insulin-like peptides in growth control. Curr. Biol. 11: 213–221.

Broughton, S. J., Piper, M. D. W., Ikeya T., Bass, T. M., Jacobson, J., Driege, Y., Martinez, P., Hafen, E., Withers, D. J., Leevers, S. J., and L. Partridge. 2005. Longer lifespan, altered metabolism, and stress resistance in Drosophila from ablation of cells making insulin-like ligands. Proc. Natl. Acad. Sci. U.S.A. 102:3105–3110.

Casas-Tinto, S., Marr II, M. T., Andreu, P., and O. Puig. 2007. Characterization of the *Drosophila* insulin receptor promoter. Biochim. Biophys. Acta 1769:236–243.

Catalán, A., Glaser-Schmitt, A., Argyridou, E., Duchen, P., and J. Parsch. 2016. An Indel Polymorphism in the *MtnA* 3’ Untranslated Region Is Associated with Gene Expression Variation and Local Adaptation in *Drosophila melanogaster*. PLoS Genet. 12:e1005987.

Chen, Y., Lee, S. F., Blanc, E., Reuter, C., Wertheim, B., Martinez-Diaz, P., Hofmann, A. A., and L. Partridge. 2012. Genome-Wide Transcription Analysis of Clinal Genetic Variation in *Drosophila*. PLoS ONE 7:e34620.

Chippindale, A. K., Leroi, A. M., Kim, S. B., and M. R. Rose. 1993. Phenotypic plasticity and selection in *Drosophila* life-history evolution. I. Nutrition and the cost of reproduction. J. Evol. Biol. 6:171–193.

Clancy, D. J., Gems, D., Harshman, L. G., Oldham, S., Stocker, H., Hafen, E., Leevers, S. J., and L. Partridge. 2001. Extension of life-span by loss of CHICO, a *Drosophila* insulin receptor substrate protein. Science 292:104–106.

Clemson, A. S., Sgrò, C. M., and M. Telonis-Scott. 2016. Thermal plasticity in *Drosophila melanogaster* populations from eastern Australia: quantitative traits to transcripts. J. Evol. Biol. 29:2447–2463.

Cogni, R., Kuczynski, K., Koury, S., Lavington, E., Behrman, E. L., O’Brien, K. R., Schmidt, P. S., and W. F. Eanes. 2017. On the Long-term Stability of Clines in Some Metabolic Genes in *Drosophila melanogaster*. Sci. Rep. 7:42766.

Cohen, J. 1988. Statistical power analysis for the behavioral sciences. 2nd edition. Lawrence Earlbaum, Hillsdale (NJ).

Conover, D. O., and E. T. Schultz. 1995. Phenotypic similarity and the evolutionary significance of countergradient variation. Trends Ecol. Evol. 10:248–252.

Cooper, B. S., Tharp II, J. M., Jernberg I. I., and M. J. Angilletta Jr. 2012. Developmental plasticity of thermal tolerances in temperate and subtropical populations of *Drosophila melanogaster*. J. Therm. Biol. 37:211–216.

Coyne, J. A., and E. Beecham. 1987. Heritability of Two Morphological Characters Within and Among Natural Populations of *Drosophila melanogaster*. Genetics 117: 727–737.

Dantzer, B., and E. M. Swanson. 2011. Mediation of vertebrate life histories via insulin-like growth factor-1. Biol. Rev. 87:414–429.

David, J. R., and C. Bocquet. 1975. Evolution in a cosmopolitan species: genetic latitudinal clines in *Drosophila melanogaster* wild populations. Experientia 31:164–166.

David, J. R., and P. Capy. 1988. Genetic variation of *Drosophila melanogaster* natural populations. Trends Genet. 4:106–111.

David, J. R., Moreteau, B., Gauthier, J. P., Pétavy, G., Stockel, A., and A. G. Imasheva. 1994. Reaction Norms of Size Characters in Relation to Growth Temperature in *Drosophila melanogaster* - an Isofemale Lines Analysis. Genet. Sel. Evol. 26:229–251.

de Jong, G., and Z. Bochdanovits. 2003. Latitudinal clines in *Drosophila melanogaster*: body size, allozyme frequencies, inversion frequencies, and the insulin-signalling pathway. J. Genet. 82: 207–223.

Duchen, P., Zivkovic, D., Hutter, S., Stephan, W., and S. Laurent. 2013. Demographic inference reveals African and European admixture in the North American *Drosophila melanogaster* population. Genetics 193:291–301.

Durmaz, E., Benson, C., Kapun, M., Schmidt, P., and T. Flatt. 2018. An Inversion Supergene in *Drosophila* Underpins Latitudinal Clines in Survival Traits. J. Evol. Biol. 31:1354–1364.

Endler, J. A. 1977. Geographic Variation, Speciation and Clines. Princeton Univ. Press, Princeton, NJ.

Fabian, D. K., Kapun, M., Nolte, V., Kofler, R., Schmidt, P. S., Schlötterer, C., and T. Flatt. 2012. Genome-wide patterns of latitudinal differentiation among populations of *Drosophila melanogaster* from North America. Mol. Ecol. 21:4748–4769.

Fabian, D. K., Lack, J. B., Mathur, V., Schlötterer, C., Schmidt, P. S., Pool, J. E., and T. Flatt. 2015. Spatially varying selection shapes life-history clines among populations of *Drosophila melanogaster* from sub-Saharan Africa. J. Evol. Biol. 28:826–840.

Fabian, D.K., Garschall, K., Klepsatel, P., Santos-Matos, G., Sucena, E., Kapun, M., Lemaitre, B., Schlötterer, C., Arking, R., and T. Flatt. 2018. Evolution of longevity improves immunity in *Drosophila*. Evol. Lett. 2:567–579.

Fielenbach, N., and A. Antebi. 2008. *C. elegans* dauer formation and the molecular basis of plasticity. Genes Dev. 22:2149–2165.

Finch, C. E., and M. R. Rose. 1995. Hormones and the physiological architecture of life history evolution. Quart. Rev. Biol. 70:1–52.

Flachsbart, F., Caliebe, A., Kleindorp, R., Blanché, H., von Eller-Eberstein, H., Nikolaus, S., Schreiber, S., and A. Nebel. 2009. Association of FOXO3A variation with human longevity confirmed in German centenarians. Proc. Natl. Acad. Sci. U.S.A. 106:2700–2705.

Flatt, T. 2004. Assessing natural variation in genes affecting *Drosophila* lifespan. Mech. Ageing Dev. 125:155–159.

Flatt, T. 2016. Genomics of clinal variation in *Drosophila*: disentangling the interactions of selection and demography. Mol. Ecol. 25:1023–1026.

Flatt, T., and A. Heyland. 2011. Mechanisms of Life History Evolution. Oxford University Press, Oxford.

Flatt, T., and L. Partridge. 2018. Horizons in the evolution of aging. BMC Biol. 16(1):93.

Flatt, T., and D. E. L. Promislow. 2007. Still pondering an age-old question. Science 318:1255–1256.

Flatt, T., and P. S. Schmidt. 2009. Integrating evolutionary and molecular genetics of aging. Biochim. Biophys. Acta 1790:951–962.

Flatt, T., Tu, M.-P., and M. Tatar. 2005. Hormonal pleiotropy and the juvenile hormone regulation of *Drosophila* development and life history. BioEssays 27:999–1010.

Flatt, T., Amdam, G. V., Kirkwood, T. B. L., and S. W. Omholt. 2013. Life-History Evolution and the Polyphenic Regulation of Somatic Maintenance and Survival. Quart. Rev. Biol. 88:185–218.

Folguera, G., Ceballos, S., Spezzi, L., Fanara, J. J., and E. Hasson. 2008. Clinal variation in developmental time and viability, and the response to thermal treatments in two species of *Drosophila*. Biol. J. Linnean Soc. 95:233–245.

Frazier, M. R., Harrison, J. F., Kirkton, S. D., and S. P. Roberts. 2008. Cold rearing improves cold-flight performance in Drosophila via changes in wing morphology. J. Exp. Biol. 211: 2116–2122.

Gems, D., Sutton, A. J., Sundermeyer, M. L., Albert, P. S., King, K. V., Edgley, M. L., Larsen, P. L., and D. L. Riddle. 1998. Two pleiotropic classes of *daf-2* mutation affect larval arrest, adult behavior, reproduction and longevity in *Caenorhabditis elegans*. Genetics 150:129–155.

Giannakou, M. E., and L. Partridge. 2007. Role of insulin-like signalling in *Drosophila* lifespan. Trends Biochem. Sci. 32:180–188.

Giannakou, M. E., Goss, M., Jünger, M. A., Hafen, E., Leevers, S. J., and L. Partridge. 2004. Long-lived *Drosophila* with overexpressed dFOXO in adult fat body. Science 305:361.

Giannakou, M. E., Goss, M., and L. Partridge. 2008. Role of dFOXO in lifespan extension by dietary restriction in *Drosophila melanogaster*: not required, but its activity modulates the response. Aging Cell 7:187–198.

Gilchrist, A. S., Azevedo, R. B. R., Partridge, L., and P. O’Higgins. 2000. Adaptation and constraint in the evolution of *Drosophila melanogaster* wing shape. Evol. Dev. 2:114–124

Goenaga, J., Fanara, J. J., and E. Hasson. 2010. A quantitative genetic study of starvation resistance at different geographic scales in natural populations of *Drosophila melanogaster*. Genet. Res. 92:253–259.

Goenaga, J., Fanara, J. J., and E. Hasson. 2013. Latitudinal Variation in Starvation Resistance is Explained by Lipid Content in Natural Populations of *Drosophila melanogaster*. Evol. Biol. 40:601–612.

Hoffmann, A. A., and L. G. Harshman. 1999. Desiccation and starvation resistance in *Drosophila*: patterns of variation at the species, population and intrapopulation levels. Heredity 83:637–643.

Hoffmann, A. A., and S. W. McKechnie. 1991. Heritable Variation in Resource Utilization and Response in a Winery Population of *Drosophila melanogaster*. Evolution 45:1000–1015.

Hoffmann, A. A., and A. R. Weeks. 2007. Climatic selection on genes and traits after a 100 year-old invasion: a critical look at the temperate-tropical clines in *Drosophila melanogaster* from eastern Australia. Genetica 129:133–147.

Hoffmann, A. A., Anderson, A., and R. Hallas. 2002. Opposing clines for high and low temperature resistance in *Drosophila melanogaster*. Ecol. Lett. 5:614–618.

Hoffmann, A. A., Shirriffs, J., and M. Scott. 2005. Relative importance of plastic vs genetic factors in adaptive differentiation: geographical variation for stress resistance in *Drosophila melanogaster* from eastern Australia. Func. Ecol. 19:222–227.

Holzenberger, M., Dupont, J., Ducos, B., Leneuve, P., Géloën, A., Even, P. C., Cervera, P., and Y. Le Bouc. 2003. IGF-1 receptor regulates lifespan and resistance to oxidative stress in mice. Nature 421:182–187.

Houle, D. 2001. Characters as the Units of Evolutionary Change. Pp. 109–140 in G. P. Wagner, ed. The Character Concept in Evolutionary Biology. Academic Press, San Diego, CA.

Hwangbo, D. S., Gersham, B., Tu, M.-P., Palmer, M., and M. Tatar. 2004. *Drosophila* dFOXO controls lifespan and regulates insulin signalling in brain and fat body. Nature 429:562–566.

James, A. C., and L. Partridge. 1995. Thermal evolution of rate of larval development in *Drosophila melanogaster* in laboratory and field populations. J. Evol. Biol. 8:315–330.

James, A. C., Azevedo, R. B., and L. Partridge. 1997. Genetic and environmental responses to temperature of *Drosophila melanogaster* from a latitudinal cline. Genetics 146:881–890.

Johnston, S. E., Gratten, J., Berenos, C., Pilkington, J. G., Clutton-Brock, T. H., Pemberton, J. M., and J. Slate. 2013. Life history trade-offs at a single locus maintain sexually selected genetic variation. Nature 502:93–95.

Jones, F. C., Grabherr, M. G., Chan, Y. F., Russell, P., Mauceli, E., Johnson, J., Swofford, R., Pirun, M., Zody, M. C., White, S., Birney, E., Searle, S., Schmutz, J., Grimwood, J., Dickson, M. C., Myers, R. M., Miller, C. T., Summers, B. R., Knecht, A. K., Brady, S. D., Zhang, H., Pollen, A. A., Howes, T., Amemiya, C., Broad Institute Genome Sequencing Platform & Whole Genome Assembly Team, Baldwin, J., Bloom, T., Jaffe, D. B., Nicol, R., Wilkinson, J., Lander, E. S., Di Palma, F., Lindblad-Toh, K., and D. M. Kingsley. 2012. The genomic basis of adaptive evolution in threespine sticklebacks. Nature 484:55–61.

Jovelin, R., Comstock, J. S., Cutter, A. D., and P.C. Phillips. 2014. A Recent Global Selective Sweep on the *age-1* Phosphatidylinositol 3-OH Kinase Regulator of the Insulin-Like Signaling Pathway Within *Caenorhabditis remanei*. G3 (Bethesda) 4:1123–1133.

Jünger, M. A., Rintelen, F., Stocker, H., Wasserman, J. D., Végh, M., Radimerski, T., Greenberg, M. E., and E. Hafen. 2003. The *Drosophila* forkhead transcription factor FOXO mediates the reduction in cell number associated with reduced insulin signaling. J. Biol. 2(3):20.

Kao, J. Y., Zubair, A., Salomon, M. P., Nuzhdin, S. V., and D. Campo. 2015. Population genomic analysis uncovers African and European admixture in *Drosophila melanogaster* populations from the south-eastern United States and Caribbean Islands. Mol. Ecol. 24:1499–1509.

Kapun, M., Schmidt, C., Durmaz, E., Schmidt, P. S., and T. Flatt. 2016a. Parallel effects of the inversion *In(3R)Payne* on body size across the North American and Australian clines in *Drosophila melanogaster*. J. Evol. Biol. 29:1059–1072.

Kapun, M., Fabian, D. K., Goudet, J., and T. Flatt. 2016b. Genomic Evidence for Adaptive Inversion Clines in *Drosophila melanogaster*. Mol. Biol. Evol. 33:1317–1336.

Karan, D., Dahiya, N., Munjal, A. K., Gibert, P., Moreteau, B., Parkash, R., and J. R. David. 1998. Desiccation and Starvation Tolerance of Adult *Drosophila*: Opposite Latitudinal Clines in Natural Populations of Three Different Species. Evolution 52:825–831.

Keller, A. 2007. *Drosophila melanogaster*’s history as a human commensal. Curr. Biol. 17:R77–R81.

Kenyon, C. 2001. A conserved regulatory system for aging. Cell 105:165–168.

Kenyon, C. J. 2010. The genetics of ageing. Nature 464:504–512.

Kenyon, C., Chang, J., Gensch, E., Rudner, A., and R. Tabtiang. 1993. A *C. elegans* mutant that lives twice as long as wild type. Nature 366:461–464.

Klepsatel, P., Gáliková, M., De Maio, N., Huber, C. D., Schlötterer, C., and T. Flatt. 2013. Variation in Thermal Performance and Reaction Norms Among Populations of *Drosophila melanogaster*. Evolution 67:3573–3587.

Klepsatel, P., Gáliková, M., Huber, C. D., and T. Flatt. 2014. Similarities and differences in altitudinal versus latitudinal variation for morphological traits in *Drosophila melanogaster*. Evolution 68:1385–1398.

Kolaczkowski, B., Kern, A. D., Holloway, A. K., and D. J. Begun. 2011. Genomic differentiation between temperate and tropical Australian populations of *Drosophila melanogaster*. Genetics 187:245–260.

Kramer, J. M., Davidge, J. T., Lockyer, J. M., and B. E. Staveley. 2003. Expression of *Drosophila* FOXO regulates growth and can phenocopy starvation. BMC Dev. Biol. 3:5.

Kramer, J. M., Slade, J. D., and B. E. Staveley. 2008. *foxo* is required for resistance to amino acid starvation in Drosophila. Genome 51:668–672.

Kubrak, O. I., Kučerová, L., Theopold, U., and D. R. Nässel. 2014. The Sleeping Beauty: How Reproductive Diapause Affects Hormone Signaling, Metabolism, Immune Response and Somatic Maintenance in *Drosophila melanogaster*. PLoS ONE 9:e113051.

Lachaise, D., Cariou, M.-L., David J. R., Lemeunier, F., Tsacas, L., and M. Ashburner. 1988. Historical biogeography of the *Drosophila melanogaster* species subgroup. Evol. Biol. 22:159–225.

Lafuente, E., Duneau, D., and P. Beldade. 2018. Genetic basis of thermal plasticity variation in *Drosophila melanogaster* body size. PLOS Genetics 14: e1007686.

Lee, K. P., and T. Jang. 2014. Exploring the nutritional basis of starvation resistance in *Drosophila melanogaster*. Func. Ecol. 28:1144–1155.

Lee, S. F., Eyre-Walker, Y. C., Rane, R. V., Reuter, C., Vinti, G., Rako, L., Partridge, L., and A. A. Hoffmann. 2013. Polymorphism in the *neurofibromin* gene, *Nf1*, is associated with antagonistic selection on wing size and development time in *Drosophila melanogaster*. Mol. Ecol. 22:2716–2725.

Levine, M. T., Eckert, M. L., and D. J. Begun. 2011. Whole-Genome Expression Plasticity across Tropical and Temperate *Drosophila melanogaster* Populations from Eastern Australia. Mol. Biol. Evol. 28:249–256.

Levins, R. 1968. Evolution in Changing in Environments. Princeton Univ. Press, Princeton, NJ.

Levins, R. 1969. Thermal Acclimation and Heat Resistance in *Drosophila* Species. Am. Nat. 103:483–499.

Li, Q., and Z. Gong. 2015. Cold-sensing regulates *Drosophila* growth through insulin-producing cells. Nat. Comm. 6:10083.

Libina, N., Berman, J. R., and C. Kenyon. 2003. Tissue-specific activities of *C. elegans* DAF-16 in the regulation of lifespan. Cell 115:489–502.

Lihoreau, M., Poissonnier, L.-A., Isabel, G., and A. Dussutour. 2016. *Drosophila* females trade off good nutrition with high-quality oviposition sites when choosing foods. J. Exp. Biol. 219:2514–2524.

Machado, H. E., Bergland, A. O., O’Brien, K. R., Behrman, E. L., Schmidt, P. S., and D. A. Petrov. 2016. Comparative population genomics of latitudinal variation in *Drosophila simulans* and *Drosophila melanogaster*. Mol. Ecol. 25:723–740.

Machado, H. E., Bergland, A. O., Taylor, R., Tilk, S., Behrman, E. L., Dyer, K., Fabian, D. K., Flatt, T., González, J., Karasov, T.L., Kozeretska, I., Lazzaro, B. P., Merritt, T. J. S., Pool, J. E., O’Brien, K., Rajpurohit, S., Roy, P. R., Schaeffer, S. W., Serga, S., Schmidt, P., and D. Petrov. 2018. Broad geographic sampling reveals predictable and pervasive seasonal adaptation in *Drosophila*. bioRxiv doi: https://doi.org/10.1101/337543.

Mackay, T. F., Stone, E. A., and J. F. Ayroles. 2009. The genetics of quantitative traits: challenges and prospects. Nat. Rev. Genet. 10:565–577.

Mackay, T. F. C., Richards, S., Stone E. A., Barbadilla, A., Ayroles, J. F., Zhu, D., Casillas, S., Han, Y., Magwire, M. M., Cridland, J. M., Richardson, M. F., Anholt, R. R., Barrón, M., Bess, C., Blankenburg, K. P., Carbone, M. A., Castellano, D., Chaboub, L., Duncan, L., Harris, Z., Javaid, M., Jayaseelan, J. C., Jhangiani, S. N., Jordan, K. W., Lara, F., Lawrence, F., Lee, S. L., Librado, P., Linheiro, R. S., Lyman, R. F., Mackey, A. J., Munidasa, M., Muzny, D. M., Nazareth, L., Newsham, I., Perales, L., Pu, L. L., Qu, C., Ràmia, M., Reid, J. G., Rollmann, S. M., Rozas, J., Saada, N., Turlapati, L., Worley, K. C., Wu, Y. Q., Yamamoto, A., Zhu, Y., Bergman, C. M., Thornton, K. R., Mittelman, D., and R. A. Gibbs. 2012. The *Drosophila melanogaster* Genetic Reference Panel. Nature 482:173–178.

Marcil, J., Swain, D. P., and J. A. Hutchings. 2006. Countergradient variation in body shape between two populations of Atlantic cod (*Gadus morhua*). Proc. Roy. Soc. London B 273:217–223.

Markow, T. A., Raphael, B., Dobberfuhl, D., Breitmeyer, C. M., Elser, J. J., and E. Pfeiler. 1999. Elemental stoichiometry of *Drosophila* and their hosts. Func. Ecol. 13:78–84.

Mathur, V., and P.S. Schmidt. 2017. Adaptive patterns of phenotypic plasticity in laboratory and field environments in *Drosophila melanogaster*. Evolution 71:465–474.

Mattila, J., Bremer, A., Ahonen, L., Kostiainen, R., and O. Puig. 2009. *Drosophila* FoxO Regulates Organism Size and Stress Resistance through an Adenylate Cyclase. Mol. Cell Biol. 29:5357–5365.

Matzkin, L. M., Watts, T. D., and T. A. Markow. 2009. Evolution of stress resistance in *Drosophila*: interspecific variation in tolerance to desiccation and starvation. Func. Ecol. 23:521–527.

McGaugh, S. E., Bronikowski, A. M., Kuo, C.-H., Reding, D. M., Addis, E. A., Flagel, L. E., Janzen, F. J., and T. S. Schwartz. 2015. Rapid molecular evolution across amniotes of the IIS/TOR network. Proc. Natl. Acad. Sci. U.S.A. 112:7055–7060.

McKechnie, S. W., Blacket, M. J., Song, S. V., Rako, L., Carroll, X., Johnson, T. K., Jensen, L. T., Lee, S. F., Wee, C. W., and A. A. Hoffmann. 2010. A clinally varying promoter polymorphism associated with adaptive variation in wing size in *Drosophila*. Mol. Ecol. 19:775–784.

Méndez-Vigo, B., Martínez-Zapater, J. M., and C. Alonso-Blanco. 2013. The Flowering Repressor *SVP* Underlies a Novel *Arabidopsis thaliana* QTL Interacting with the Genetic Background. PLoS Genet. 9:e1003289.

Murphy, C. T., McCarroll, S. A., Bargmann, C. I., Frasser, A., Kamath, R. S., Ahringer, J., Li, H., and C. Kenyon. 2003. Genes that act downstream of DAF-16 to influence the lifespan of *Caenorhabditis elegans*. Nature 424:277–283.

Oldham, S., and E. Hafen. 2003. Insulin/IGF and target of rapamycin signaling: a TOR de force in growth control. Trends Cell Biol. 13:79–85.

Oldham, S., Stocker, H., Laffargue, M., Wittwer, F., Wymann, M., and E. Hafen. 2002. The *Drosophila* insulin/IGF receptor controls growth and size by modulating PtdIns P3 levels. Development 129:4103–4109.

Overgaard, J., Kristensen, T. N., Mitchell, K. A., and A. A. Hoffmann. 2011. Thermal Tolerance in Widespread and Tropical *Drosophila* Species: Does Phenotypic Plasticity Increase with Latitude? Am. Nat. 178:S80–S96.

Paaby, A. B., and P. S. Schmidt. 2009. Dissecting the genetics of longevity in *Drosophila melanogaster*. Fly (Austin) 3:1–10.

Paaby, A. B., Bergland, A. O., Behrman, E. L., and P. S. Schmidt. 2014. A highly pleiotropic amino acid polymorphism in the *Drosophila* insulin receptor contributes to life-history adaptation. Evolution 68:3395–3409.

Paaby, A. B., Blacket, M. J., Hoffmann, A. A., and P. S. Schmidt. 2010. Identification of a candidate adaptive polymorphism for *Drosophila* life history by parallel independent clines on two continents. Mol. Ecol. 19:760–774.

Partridge, L., and D. Gems. 2002. Mechanisms of ageing: public or private? Nat. Rev. Genet. 3:165–175.

Partridge, L., Barrie, B., Fowler, K., and V. French. 1994a. Evolution and development of body size and cell size in *Drosophila melanogaster* in response to temperature. Evolution 48:1269–1276.

Partridge, L., Barrie, B., Fowler, K., and V. French, V. 1994b. Thermal Evolution of Pre-Adult Life-History Traits in *Drosophila melanogaster*. J. Evol. Biol. 7:645–663.

Partridge, L., Gems, D., and D. J. Withers, D. J. 2005. Sex and Death: What Is the Connection? Cell 120:461–472.

Ponton, F., Chapuis, M.-P., Pernice, M., Sword, G. A., and S. J. Simpson. 2011. Evaluation of potential reference genes for reverse transcription-qPCR studies of physiological responses in *Drosophila melanogaster*. J. Insect Physiol. 57:840–850.

Puig, O., and J. Mattila. 2011. Understanding Forkhead Box Class O Function: Lessons from *Drosophila melanogaster*. Antiox. Redox Sign. 14:635–647.

Puig, O., and R. Tjian. 2005. Transcriptional feedback control of insulin receptor by dFOXO / FOXO1. Genes Dev. 19:2435–2446.

Puig, O., Marr, M. T., Ruhf, M. L., and R. Tjian. 2003. Control of cell number by *Drosophila* FOXO: downstream and feedback regulation of the insulin receptor pathway. Genes Dev. 17:2006–2020.

Rajpurohit, S., Gefen, E., Bergland, A.O., Petrov, D.A., Gibbs, A.G., and P. S. Schmidt. 2018. Spatiotemporal dynamics and genome-wide association genome-wide association analysis of desiccation tolerance in *Drosophila melanogaster*. Mol. Ecol. 27:3525–3540.

Reinhardt, J. A., Kolaczkowski, B., Jones, C. D., Begun, D. J., and A. D. Kern. 2014. Parallel Geographic Variation in *Drosophila melanogaster*. Genetics 197:361–373.

Reis, T. 2016. Effects of Synthetic Diets Enriched in Specific Nutrients on *Drosophila* Development, Body Fat, and Lifespan. PLoS ONE 11:e0146758.

Remolina, S.C., Chang, P.L., Leips, J., Nuzhdin, S.V., and K. A. Hughes. 2012. Genomic Basis of Aging and Life History Evolution in *Drosophila melanogaster*. Evolution 66:3390–3403.

Reznick, D. N. 2005. The genetic basis of aging: an evolutionary biologist’s perspective. Sci. Aging Knowl. Env. (SAGE KE) 11:pe7.

Robinson, S. J. W., Zwaan, B., and L. Partridge. 2000. Starvation Resistance and Adult Body Composition in a Latitudinal Cline of *Drosophila melanogaster*. Evolution 54:1819–1824.

Rockman, M. V. 2012. The QTN program and the alleles that matter for evolution: all that’s gold does not glitter. Evolution 66:1–17.

Sawilowsky, S. 2009. New effect size rules of thumb. J. Mod. Appl. Stat. Meth. 8:467–474.

Schiesari, L., Andreatta, G., Kyriacou, C. P., O’Connor, M. B., and R. Costa. 2016. The Insulin-Like Proteins dILPs-2/5 Determine Diapause Inducibility in *Drosophila*. PLoS ONE 11:e0163680.

Schluter, D., Price, T. D., and L. Rowe. 1991. Conflicting selection pressures and life history trade-offs. Proc. Roy. Soc. London B 246:11–17.

Schmidt, P. S., and D. R. Conde. 2006. Environmental Heterogeneity and the Maintenance of Genetic Variation for Reproductive Diapause in *Drosophila melanogaster*. Evolution 60:1602–1611.

Schmidt, P. S., and A. B. Paaby. 2008. Reproductive Diapause and Life-History Clines in North American Populations of *Drosophila melanogaster*. Evolution 62:1204–1215.

Schmidt, P. S., Duvernell, D. D., and W. F. Eanes. 2000. Adaptive evolution of a candidate gene for aging in *Drosophila*. Proc. Natl. Acad. Sci. U.S.A. 97:10861–10865.

Schmidt, P. S., Matzkin, L., Ippolito, M., and W. F. Eanes. 2005a. Geographic Variation in Diapause Incidence, Life-History Traits, and Climatic Adaptation in *Drosophila melanogaster*. Evolution 59:1721–1732.

Schmidt, P. S., Paaby, A. B., and M. S. Heschel. 2005b. Genetic variance for diapause expression and associated life histories in *Drosophila melanogaster*. Evolution 59:2616–2625.

Schmidt, P. S., Zhu, C-T., Das, J., Batavia, M., Yang, L., and W. F. Eanes. 2008. An amino acid polymorphism in the *couch potato* gene forms the basis for climatic adaptation in *Drosophila melanogaster*. Proc. Natl. Acad. Sci. U.S.A. 105:16207–16211.

Schwartz, T. S., and A. M. Bronikowski. 2016. Evolution and Function of the Insulin and Insulin-like Signaling Network in Ectothermic Reptiles: Some Answers and More Questions. Integr. Comp. Biol. 56:171–184.

Sgrò, C. M., Overgaard, J., Kristensen, T. N., Mitchell, K. A., Cockerell, F. E., and A. A. Hoffmann. 2010. A comprehensive assessment of geographic variation in heat tolerance and hardening capacity in populations of *Drosophila melanogaster* from eastern Australia. J. Evol. Biol. 23: 2484–2493.

Siddiq, M. A., Loehlin, D. W., Montooth, K. L., and J. W. Thornton. 2017. Experimental test and refutation of a classic case of molecular adaptation in *Drosophila melanogaster*. Nat. Ecol. Evol. 1(2):0025.

Slack, C., Giannakou, M. E., Foley, A., Goss, M., and L. Partridge. 2011. dFOXO-independent effects of reduced insulin-like signaling in *Drosophila*. Aging Cell 10: 735–748.

Sparkman, A. M., Vleck, C. M., and A. M. Bronikowski. 2009. Evolutionary ecology of endocrine-mediated life-history variation in the garter snake *Thamnophis elegans*. Ecology 90:720–728.

Sparkman, A. M., Byars, D., Ford, N. B., and A. M. Bronikowski. 2010. The role of insulin-like growth factor-1 (IGF-1) in growth and reproduction in female brown house snakes (*Lamprophis fuliginosus*). Gen. Comp. Endocrinol. 168:408–414.

Stalker, H. D. 1980. Chromosome-Studies in Wild Populations of *Drosophila melanogaster*. II. Relationship of Inversion Frequencies to Latitude, Season, Wing-Loading and Flight Activity. Genetics 95(1):211–223.

Stern, D. L. 2000. Perspective: evolutionary developmental biology and the problem of variation. Evolution 54:1079–1091.

Stern, D. L., 2011. Evolution, Development, & the Predictable Genome. Roberts & Co. Publishers, Greenwood Village, CO.

Strassburger, K., Zoeller, T., Sandmann, T., Leible, S., Kerr, G., Boutros, M., and A. A. Teleman. 2017. Sorting & Sequencing Flies by Size: Identification of Novel TOR Regulators and Parameters for Successful Sorting. bioRxiv doi: https://doi.org/10.1101/119719

Stuart, J. A., and M. M. Page. 2010. Plasma IGF-1 is negatively correlated with body mass in a comparison of 36 mammalian species. Mech. Ageing Dev. 131:591–598.

Suh, Y., Atzmon, G., Cho, M.-O., Hwang, D., Liu, B., Leahy, D. J., Barzilai, N, and P. Cohen. 2008. Functionally significant insulin-like growth factor I receptor mutations in centenarians. Proc. Natl. Acad. Sci. U.S.A. 105:3438–3442.

Svetec, N., Saelao, P., Cridland, J. M., Hoffmann, A. A., and D. J. Begun. 2018. Functional Analysis of a Putative Target of Spatially Varying Selection in the *Menin1* Gene of *Drosophila melanogaster*. G3 (Bethesda), in press.

Swanson, E. M., and B. Dantzer. 2014. Insulin-like growth factor-1 is associated with life-history variation across Mammalia. Proc. Roy. Soc. London B 281:20132458.

Tang, H. Y., Smith-Caldas, M. S. B., Driscoll, M. V., Salhadar, S., and A. W. Shingleton. 2011. FOXO regulates organ-specific phenotypic plasticity in *Drosophila*. PLoS Genet. 7(11):e1002373.

Tatar, M., and C.-M. Yin. 2001. Slow aging during insect reproductive diapause: why butterflies, grasshoppers and flies are like forms. Exp. Gerontol. 36:723–738.

Tatar, M., Bartke, A., and A. Antebi. 2003. The Endocrine Regulation of Aging by Insulin-like Signals. Science 299:1346–1351.

Tatar, M., Kopelman, A., Epstein, D., Tu, M.-P., Yin, C.-M., and R. S. Garofalo. 2001. A mutant *Drosophila* insulin receptor homolog that extends life-span and impairs neuroendocrine function. Science 292:107–110.

Teleman, A. A. 2010. Molecular mechanisms of metabolic regulation by insulin in *Drosophila*. Biochem. J. 425:13–26.

Tennessen, J. M., Barry, W. E., Cox, J., and C. S. Thummel. 2014. Methods for studying metabolism in *Drosophila*. Methods 68:105–115.

Trotta, V., Calboli, F. C. F., Ziosi, M., Guerra, D., Pezzoli, M. C., David, J. R., and S. Cavicchi. 2006. Thermal plasticity in *Drosophila melanogaster*: A comparison of geographic populations. BMC Evol. Biol. 6:67.

Turner, T. L. 2014. Fine-mapping natural alleles: quantitative complementation to the rescue. Mol. Ecol. 23:2377–2382.

Turner, T. L., Levine, M. T., Eckert, M. L., and D. J. Begun. 2008. Genomic Analysis of Adaptive Differentiation in *Drosophila melanogaster*. Genetics 179:455–473.

van Heerwaarden, B., and C. M. Sgrò. 2017. The quantitative genetic basis of clinal divergence in phenotypic plasticity. Evolution 71:2618–2633.

van ‘t Land, J., van Putten, P., Zwaan, B., Kamping A., and W. van Delden. 1999. Latitudinal variation in wild populations of *Drosophila melanogaster*: heritabilities and reaction norms. J. Evol. Biol. 12:222–232.

Vonesch, S. C., Lamparter, D., Mackay, T. F. C., Bergmann, S., and E. Hafen. 2016. Genome-Wide Analysis Reveals Novel Regulators of Growth in *Drosophila melanogaster*. PLoS Genet. 12:e1005616.

Wang, K., Dickson, S. P., Stolle, C. A., Krantz, I. D., Goldstein, D. B., and H. Hakonarson. 2010. Interpretation of Association Signals and Identification of Causal Variants from Genome-wide Association Studies. Am. J. Hum. Genet. 86:730–742.

Weeks, A. R., McKechnie, S. W., and A. A. Hoffmann. 2002. Dissecting adaptive clinal variation: markers, inversions and size/stress associations in *Drosophila melanogaster* from a central field population. Ecol. Lett. 5:756–763.

Whitlock, M. C., and D. Schluter. 2009. The Analysis of Biological Data. Roberts and Company Publishers, Greenwood Village, CO.

Willcox, B. J., Donlon, T. A., He, Q., Chen, R., Grove, J. S., Yano, K., Masaki, K. H., Willcox, D. C., Rodriguez, B., and J. D. Curb. 2008. FOXO3A genotype is strongly associated with human longevity. Proc. Natl. Acad. Sci. U.S.A. 105:13987–1399.

Williams, G. C. 1957. Pleiotropy, natural selection, and the evolution of senescence. Evolution 11:398–411.

Williams, K. D., Busto, M., Suster, M. L., So, M. L., So, A. K.-C., Ben-Shahar, Y., Leevers, S. J., and M. B. Sokolowski. 2006. Natural variation in *Drosophila melanogaster* diapause due to the insulin-regulated PI3-kinase. Proc. Natl. Acad. Sci. U.S.A. 103:15911–15915.

Williams, K. D., and M. B. Sokolowski. 1993. Diapause in *Drosophila melanogaster* females: a genetic analysis. Heredity 71:312–317.

Zhang, B., Xiao, R., Ronan, E. A., He, Y., Hsu, A-L., Liu, J., and X. Z. Xu. 2015. Environmental Temperature Differentially Modulates *C. elegans* Longevity through a Thermosensitive TRP Channel. Cell Rep. 11:1414–1424.

Zhao, L., Wit, J., Svetec, N., and D. J. Begun. 2015. Parallel Gene Expression Differences between Low and High Latitude Populations of *Drosophila melanogaster* and *D. simulans*. PLoS Genet. 11:e1005184.

Zhao, X., Bergland, A. O., Behrman, E. L., Gregory, B. D., Petrov, D. A., and P.S. Schmidt. 2016. Global Transcriptional Profiling of Diapause and Climatic Adaptation in *Drosophila melanogaster*. Mol. Biol. Evol. 33:707–720.

